# Parallel processing of sensory cue and spatial information in the Dentate Gyrus

**DOI:** 10.1101/2020.02.13.947903

**Authors:** Sebnem N. Tuncdemir, Andres D. Grosmark, Gergely F. Turi, Amei Shank, Jack Bowler, Gokhan Ordek, Attila Losonczy, Rene Hen, Clay Lacefield

## Abstract

During exploration, animals form an internal map of an environment by combining information about specific sensory cues or landmarks with the animal’s motion through space, a process which critically depends on the mammalian hippocampus. The dentate gyrus (DG) is the first stage of the hippocampal trisynaptic circuit where self-motion and sensory cue information are integrated, yet it remains unknown how neurons within the DG encode both cue related (“what”) and spatial (“where”) information during cognitive map formation. Using two photon calcium imaging in head fixed mice running on a treadmill, along with on-line sensory cue manipulation at specific track locations, we have identified robust sensory cue responses in DG granule cells largely independent of spatial location. Granule cell cue responses are stable for long periods of time, selective for the modality of the stimulus and accompanied by strong inhibition of the firing of other active neurons. At the same time, there is a smaller fraction of neurons whose firing is spatially tuned but insensitive to the presentation of nearby cues, similar to traditional place cells. These results demonstrate the existence of “cue cells” in addition to the better characterized “place cells” in the DG, an important heterogeneity that has been previously overlooked. We hypothesize that the granule cell population may support multiple channels of spatial and non-spatial information that contribute distinctly to local and down-stream computations and impact the role of the dentate gyrus in spatial navigation and episodic memory.

## Introduction

An animal’s location in an environment is highly relevant for guiding its behavior, both to find areas of potential reward and avoid areas of possible danger. The mammalian hippocampal formation plays a cardinal role in navigation and spatial memory by integrating self-motion and sensory cue information into a cognitive map of an environment, exemplified by the presence of “place cells” selective for specific locations within space^1^. As the initial stage in the ‘trisynaptic circuit’^2^, the dentate gyrus (DG) is the first region in the hippocampus to integrate sensory and self-motion information into a discrete spatial representation, and thus represents the most basic state of spatial map formation^3,4^. Yet causal evidence for conjunctive encoding of sensory and spatial information by the principle neurons of the DG, granule cells, is still lacking. While recent studies have shown that granule cells may be activated by visual^5,6^, tactile^7^ and olfactory^8^ cues, their relationship to canonical ‘place cells’ and their influence on spatial map formation in the DG have not been investigated.

The DG receives its main long-range excitatory inputs from the lateral and medial entorhinal cortices (LEC and MEC, respectively), and sends mossy fiber projections only to area CA3^9^. The LEC is thought to primarily represent information about sensory cues, while the MEC is thought to more prominently encode self-motion information^10–12^. Since individual granule cell dendrites receive projections from both of these areas, these inputs have the potential to be the basis for dendritic computations that combine sensory and self-motion information into a discrete spatial representation^6,13–16^. In concert with a highly effective winner-take-all process mediated by lateral inhibition^17–19^, these conjunctive representations could facilitate behavioral discrimination of nearby salient locations with high spatial resolution^6,20,21^. Such processes may underlie proposed computational roles of the DG in hippocampal information processing such as pattern separation, where similar inputs are represented distinctly within the population in order to aid the selectivity of spatial behavior and memory^18,22^.

We sought to examine how the DG participates in spatial map formation by recording calcium activity in large populations of granule cells in the mouse dorsal DG during head-fixed locomotion on a treadmill. By controlling the administration of sensory cues and their pairing with the animal’s position on the treadmill, we were able to dissect sensory and spatial contributions to granule cell firing. We found that surprisingly most of the task-associated neurons were highly sensitive to specific sensory cues presented along the treadmill belt, rather than discrete locations. Cue responses in single neurons were stable for long periods of time, sensory modality-selective and led to inhibition of other active neurons. At the same time, a smaller fraction of neurons exhibited robust spatial tuning independent of local cue presentation, yet were more context selective. These two channels of information, sensory and spatial, were largely distinct within the granule cell population, and led us to postulate the existence of “cue cells” in addition to the better characterized “place cells” of the region. This work suggests that the DG maintains a largely separate code for cues in an environment and their location; yet displays specific points of integration, for example through mutual inhibition and spatial modulation of cue response amplitudes. These properties of the heterogeneous population of cue cells and place cells may play a role in higher level functions of the dentate gyrus such as pattern separation and contextual encoding in spatial navigation.

## Results

To investigate the interaction between sensory and spatial representations in the dentate gyrus (DG), we recorded the activity of large populations of granule cells using two photon calcium imaging in head-restrained mice running on a linear treadmill track^23^. Mice were injected unilaterally with rAAV(1/2).Syn.GCaMP6s to express the genetically encoded calcium indicator GCaMP6s in the dorsal DG. This was followed by the implantation of a chronic imaging window above the hippocampal fissure, which allowed us to image the calcium dynamics of hundreds of neurons in the granule cell layer simultaneously during exploratory behavior on the treadmill (Figure 1a). After habituation to head fixation, mice were trained to run in order to receive randomly delivered water rewards on a 2m-long treadmill belt, during 15 minute sessions (Figure 1b, Methods). Movies of population calcium imaging data were motion corrected offline^24^, and the activity of putative single neurons was isolated^25,26^. Neurons with significant spatial tuning of calcium activity along the track were identified using previously described methods^23,27^, and in a subset of sessions activity of individual neurons was tracked in multiple conditions over multiple sessions^28^.

**Figure1:**
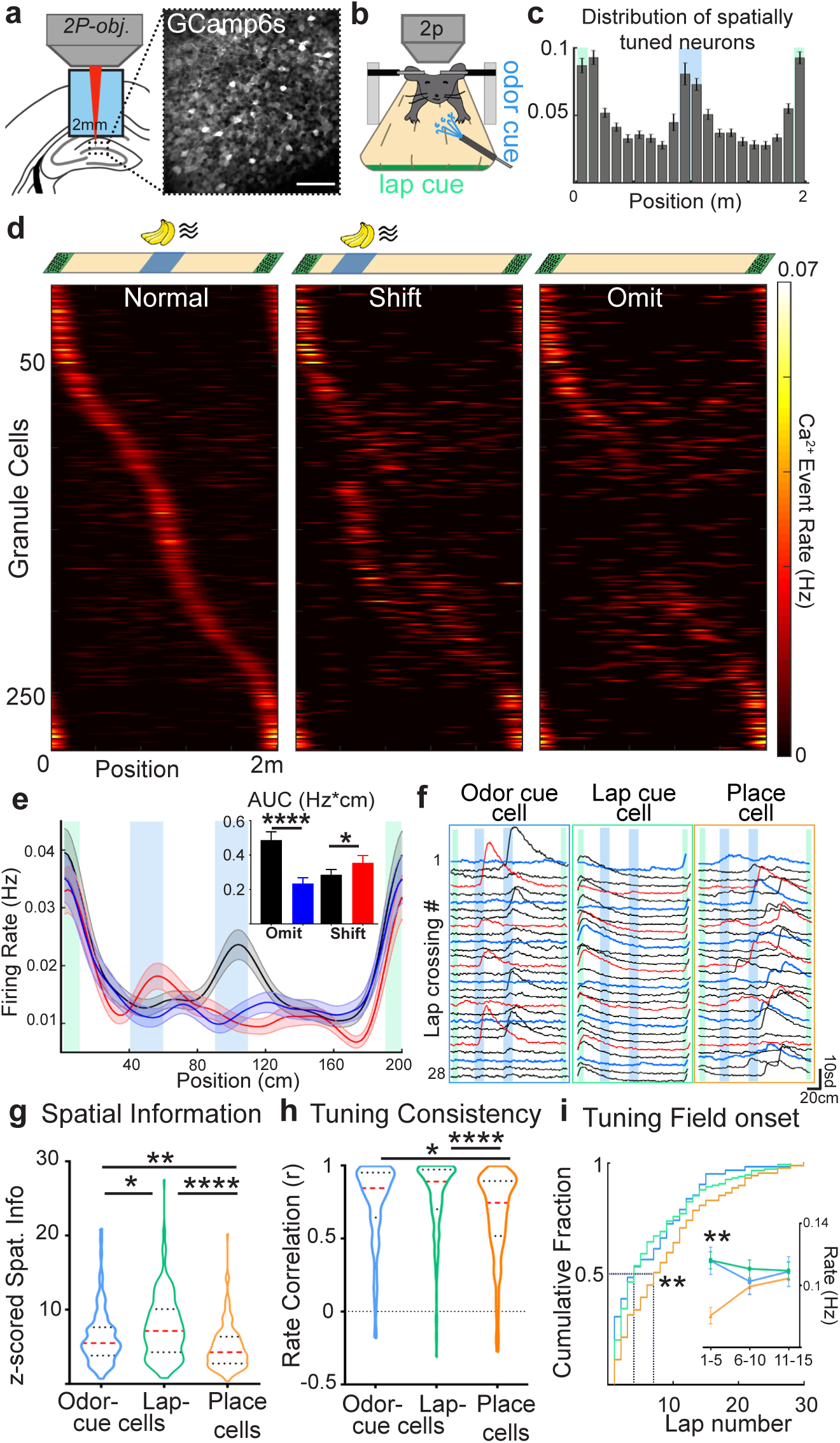
Robust representation of sensory cues in the dentate gyrus. **a)** Two-photon imaging of DG population calcium activity *in vivo*. Left, diagram of the imaging window implant in the dorsal dentate gyrus. Right, Time averaged two-photon image of GCaMP6s-expressing granule cells (Scale bar = 50μm) **b)** Diagram of head-fixed treadmill apparatus for sensory cue delivery in the spatial cue task. **c)** Fraction of spatially tuned cells at each treadmill position (bins=10cm, n = 4091 cells from 8 mice across 6-9 sessions/mouse), Locations of the odor and lap cue are shown blue and green shaded areas, respectively. **d)** DG neuron activity during the spatial cue task. Top: Location of the lap cue (green boxes) and an odor cue (blue box), on normal, and odor cue shift or omit laps. Bottom: Lap-averaged spatial firing rates of 285 spatially tuned neurons (n=8 mice) during the first session with the odor cue on normal middle location (left), cue-shifted (middle) and cue-omitted (right) laps. Each row represents activity of a single cell in the different lap types, sorted by activity on normal middle cue laps. **e)** Average spatial firing rates by position for neurons shown in panel “d.” on normal (black), cue-shifted (red) and cue-omitted laps (blue, mean ± SEM). (Inset) averaged area under the firing rate curves (AUC, Hz*cm) within the middle cue region during normal (black bar), cue shifted (red bar), or cue omitted laps (blue bar). *P_normal-omit_* <0.0001, *P_normal-shift_* =0.02, Wilcoxon signed rank test; error bars are mean ± SEM. **f)** Example fluorescence traces from representative odor-cue, lap-cue and place cells within a single session. Black, red, and blue traces represent normal, cue shifted and omitted laps, respectively. Scale bars are cm and standard deviation from each cell’s baseline fluorescence. **g)** Spatial information within cue and place cell populations. Z scored spatial information, χ^2^=48.47, p<0.0001, *P_OdorCue-LapCue_*=0.0387, *P_OdorCue-Place_*=0.0028, *P_LapCue-Place_*<0.0001, Kruskal Wallis and Dunn’s multiple comparisons test. **h)** Tuning consistency of cue and place cells. Firing rate correlation between first and last halves of the session, χ^2^=32.60, p<0.0001, *P_OdorCue-LapCue_*=0.2224, *P_OdorCue-Place_*=0.0117, *P_LapCue-Place_*<0.0001, Kruskal Wallis and Dunn’s multiple comparisons test. Red dotted lines in violin plots show median, black dotted lines show quartiles. **i)** Emergence of cue and place responses. Cumulative distribution of spatial field onset lap for cue and place cells, χ^2^=9.29, p=0.0096, *P_OdorCue-LapCue_*=0.8925, *P_OdorCue-Place_*=0.0194, *P_LapCue-Place_*=0.0091, Kruskal Wallis and Dunn’s multiple comparisons test. (Inset) Spatial field firing rates of cue and place cells during -1-5^th^, 6-10^th^ and 11-15^th^ laps, main effect of cell type: F_2,1473_=6.73, p=0.0012; cell type × lap number interaction F_4,1473_=2.44, p=0.04; main effect of lap number: F_2,1473_=0.28, p=0.7; Lap 1-5^th^, *P_OdorCue-LapCue_*=0.99, *P_OdorCue-Place_*=0.0011, *P_LapCue-Place_*<0.0001, 2-way ANOVA and Tukey’s multiple comparisons test. *N_OdorCue_*=114, *N_LapCue_*=220, *N_PlaceCell_*=160, from 8 mice 2 sessions each. \**P* < 0.05; \*\**P* < 0.01; \*\*\**P* < 0.001.

### Granule Cell Cue Responses

Following initial training on the treadmill, we introduced a 1s odor pulse delivered in the middle of the track on each lap as a dynamic spatial cue, in addition to an invariant tactile cue at the lap boundary. We found that the majority of spatially selective neurons exhibited receptive fields near the lap boundary and middle locations, corresponding to the lap cue and middle cue positions, respectively (Figure 1c, 57% of cells with peak activity within 10cm of cues, 43% >10cm from cues, p<0.0001, Mann-Whitney test). In order to normalize locomotion dependent modulation of hippocampal activity^29,30^, the treadmill was motorized at a constant speed, adjusted for each mouse (motorized velocity = 10.11 ± 0.64 cm/s, self-driven velocity = 12.76 ± 2.45 cm/s, p>0.05, Mann-Whitney test). In a separate cohort of mice advancing the treadmill belt through self-driven locomotion, we confirmed that motorizing the treadmill does not alter the representation of sensory cue information in the DG (Figures S1a-f).

Neurons responding reliably at cue locations could however be place cells that are enriched at the locations of salient stimuli, as has been suggested in area CA1^31–36^, or alternately could be directly driven by the stimulus^8^. In order to dissociate cue responses from track location, the olfactory cue was omitted or shifted 1/4^th^ of the track length (0.5m) once every 3-5 trials interleaved throughout the session (41± 2 total laps/session). Under these conditions, a majority of neurons normally active in the middle of the track shifted their firing position to match the new location of the odor in cue-shift laps (Figure 1d, middle, 79.07 ± 2.76 %) and exhibited reduced activity in cue-omitted laps (Figure 1d, right, 78.2 ± 3.87 %), compared with normal middle cue laps (n = 285 spatially tuned neurons from 8 mice). In contrast, neurons firing at locations corresponding to the invariant lap cue were unchanged in omit and shift laps. On trials with a shifted middle cue, average firing rates were higher within the new cue region (Figure 1e, red, inset p=0.0234 Wilcoxon signed-rank test), and lower at the normal cue location on cue-omitted trials (blue, inset p<0.00001, Wilcoxon signed-rank test) compared to the corresponding positions during normal trials. Rapid cue-associated activity and enrichment of highly tuned neurons at positions corresponding to sensory cues were also seen in mice during self-driven locomotion on the treadmill (Figure S1b,d). Thus, a substantial population of spatially tuned DG neurons in the virtual linear track environment are in fact active directly in response to presentation of cues at those locations, rather than the locations themselves.

Comparable population activity was found in response to cues of other sensory modalities, such as visual or tactile cues (Figures S2a-d), as well for liquid rewards (Figures S2e-h), indicating strong representation of diverse sensory variables in the DG. In addition, increasing the complexity of the environment with two additional cues at other track locations results in an additive pattern of single cue responses (Figures S2i-k), suggesting both that cue activity at different locations is relatively independent and that strong cue responses are not limited to situations with a single cue alone. Cue responses were also similar for cells imaged in the dorsal DG of a granule cell-specific transgenic mouse (Dock10 Cre^37^, Figures S3a-d,), indicating that sensory cue representations are indeed a property of the granule cell population rather than arising from other local neuron types such as mossy cells or inhibitory interneurons^7^.

Next, we divided the spatially tuned population of granule cells into three groups for subsequent analyses based on the position of their spatial fields and their activity during cue manipulation (omit or shift) trials. The three response types are illustrated for one session (Figure 1f): 1) cells with spatial fields within the middle cue region that closely track the changes in cue presentation (“odor-cue cells”, left); 2) cells with spatial fields around the lap boundary cue on the treadmill belt (“lap-cue cells”, center); and 3) the remaining spatially tuned cells with receptive fields outside of the cue locations throughout the track (“place cells”, right). These three groups, odor-cue, lap-cue, and place cells, constituted 22.5 ± 2%, 47.1 ± 2% and 30.4 ± 1% of the spatially tuned cells within the imaging field of view, respectively (Figure S3e). We found that cells with similar response types did not appear to cluster together spatially within the imaging field and the groups did not exhibit significant differences in overall mean firing rates (Figures S3e-g). Spatial coding properties however differed between cue and place coding populations of granule cells. Both populations of cue cells (lap cue and olfactory cue) showed higher average spatial information content^38^ (Fig. 1g, χ^2^=48.47, p<0.0001, Kruskal-Wallis test) and had more consistent spatial firing between the first and the second half of each session than place cells (Figure 1h, χ^2^=32.60, p<0.0001, Kruskal-Wallis test).

In order to probe the emergence of selective firing in these populations, we identified the lap in which responses began to robustly occur within the preferred spatial location of each cell during the first session of exposure to the odor cue (field onset lap^39^, see Methods). We found that for odor and lap cue cells the majority had spatial fields that appeared within the first five laps (38/65 (58%), 75/136 (55%), respectively) while the majority of place cells emerged later in the session (52/84 (62%) within 10 laps, Fig. 1i, χ^2^=9.29, p=0.0096, Kruskal-Wallis test). In agreement with the later emergence of place cells, we found that the in-field firing rates of place cells were significantly smaller than those of odor and lap cue cells only within the first 5 laps (Figure 1i, inset, 2-way ANOVA, main effect of cell type: F_2,1473_=6.73, p=0.0012; cell type × lap number interaction F_4,1473_=2.44, p=0.04; main effect of lap number: F_2,1473_=0.28, p=0.7). Additional experiments also showed that on an uncued treadmill belt prior to olfactory cue sessions, the majority of ‘future’ odor cue cells were either inactive or had low spatial tuning (Figure S4), suggesting that cue cells rapidly emerge *de novo* and are not place cells that remap to new cue locations. Taken together, these results demonstrate that sensory cue representations are more reliable and appear with less exposure than place cell representations, and arise from an independent population of granule cells.

We performed two experiments in separate cohorts of mice to exclude the possibility that the cue cells are tracking the periodicity of the stable locomotion on the motorized treadmill (i.e. are instead ‘time cells’). First, we directly compared the odor cue cell responses in mice running on a motorized treadmill at a constant speed with responses from the same cells on a self-driven treadmill at variable speeds in consecutive sessions (Figure 2a). Despite notable differences in temporal occupancy of the cue location in the two conditions (Figure S1c), the same cells fired reliably to the olfactory cue. Correlations in the spatial firing rates of individual cells show that while both groups of cue cells were highly correlated between self-driven and motorized treadmill conditions, place cells were significantly less consistent in sessions with different locomotive behaviors (Figure 2b, p<0.001, Kruskall-Wallis test). Spatial population vector correlations of all significantly tuned neurons between different locomotive behaviors also confirm highly correlated activity selectively around the cue locations (Figure 2c). Second, we investigated the latency of Ca^2+^ responses in sessions where the treadmill was motorized at different velocities each lap (Figure 2c). We reasoned that if the cue cells were tracking the period from one cue delivery to the next, cue-triggered response latencies would be proportional to the time it takes for the animal to traverse a lap while position-triggered response latencies would be constant. On the contrary, we found shorter response latencies from a constant position when the treadmill is motorized at higher velocities and vice versa, with constant cue-triggered Ca^2+^ response latencies regardless of the velocity of the animal (Figures 2e-f, S1g-h). And as in our previous experiments, omitting the cue on intermittent laps still had the effect of eliminating the responses of cue cells on that lap, confirming their acute cue sensitivity. These experiments demonstrate that sensory cue responses occur independent of volitional locomotion and that cue cells are not in fact time cells. Furthermore the differing responses of cue and place cells in these conditions supports our previous findings that these are separate, dissociable population of granule cells in the DG. For the rest of our study we will continue to call the remaining group of non cue-responsive neurons “place cells”, as they are spatially tuned cells that do not respond directly to local cues, and would therefore be called place cells based upon standards used in previous work on hippocampal spatial encoding. However, our results show that spatial responses in the majority of the dentate gyrus place cells imaged in our experiments are highly sensitive to the behavioral mode as well as the velocity of the mice, as has been shown in area CA1^36,40^, suggesting that these “place cells” respond to more than simple spatial information.

**Figure 2:**
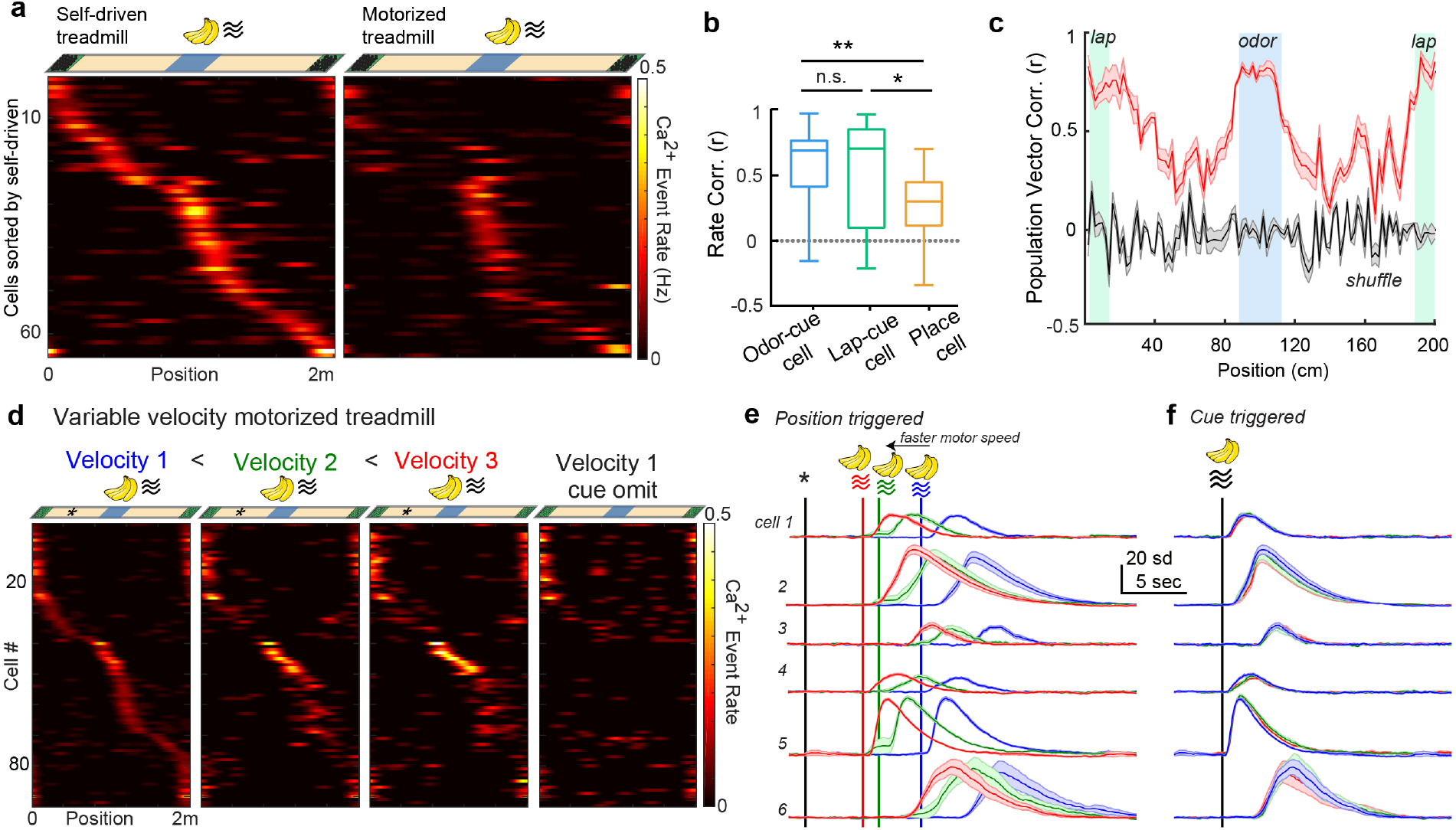
Stable encoding of sensory cues regardless of animal’s locomotive behavior or velocity. **a)** Activity of DG granule cells in response to the same odor-cued track during self-driven (left) and motorized (right) treadmill locomotion. Top: diagram of treadmill track with lap cue and middle odor cue. Bottom: Spatial firing rates for neurons tracked through consecutive sessions of self-driven and motorized treadmills (n=67 cells, 5 mice), cells are sorted by their activity in the self-driven treadmill session. See also Figure S1a-f. **b)** Comparison of average spatial rate correlations in individual neurons tracked across self-driven and motored treadmills (χ^2^=10, p<0.01, *P_Odor-Lap cue_*>0.99, *P_Odor Cue-Place_* =0.0059, *P_Lap Cue-Place_* =0.0421, *N_Odor-Cue_*=30, *N_Lap-Cue_*=22, *N_Place_*=15). Kruskal-Wallis test, Dunn’s multiple comparisons test. Boxes, 25th to 75th percentiles; bars, median; whiskers, 99% range. \**P* < 0.05; \*\**P* < 0.01. **c)** Spatial population vector correlations of spatially tuned neurons between self-driven and motorized treadmills n=31±9 cells in 5 mice (red), shaded error bars represent mean ± SEM for each treadmill position. Locations of the middle and lap cue are shown by blue and green shaded areas, respectively. Mean population vector correlations by mice exceeded chance levels as estimated by shuffling cell identity in each mouse/session (gray). **d)** DG neuron activity during the spatial cue task when the treadmill is motorized at different velocities in each lap, or the cue omitted, in a pseudorandom manner. Top: velocity1: 6cm/s, velocity2: 9cm/sec, velocity3: 12cm/sec, asterisks represent reference location for position-triggered averages in following panels; Bottom: Spatial firing rates of 89 spatially tuned neurons (n=3 mice) during different velocity and cue-omitted laps. Each row across all graphs represents a single cell, with the population sorted by each cell’s activity on velocity1 laps. See also Figure S1g-h. **e)** Example reference *position-triggered* average Ca^2+^ transients from six example cue cells on faster (red), normal (green), and slower (blue) velocity laps. Axes are seconds and standard deviation from each cell’s baseline fluorescence for the session. **f)** Example *cue-triggered* average Ca^2+^ transients from the same cue cells shown in panel “e.” on faster (red), normal (green), and slower (blue) velocity laps.

### Stability and Specificity of Sensory Cue Responses

To further characterize the stability and coding specificity among the granule cell subpopulations, we investigated the responses of individual neurons over time and with respect to different sensory cues (Figure 3a, b). We tracked cells^28^ over multiple sessions and were able to find substantial numbers of the same cells active in different sessions within a day or 1 week later in the same fields of view (Figures S5a, b). Although not all cells were identified in every session, a similar percentage of spatially selective neurons was registered in all sessions, which was confirmed by visual inspection to ensure that cells appeared consistent in the anatomical images (Figures S5c-g).

**Figure 3:**
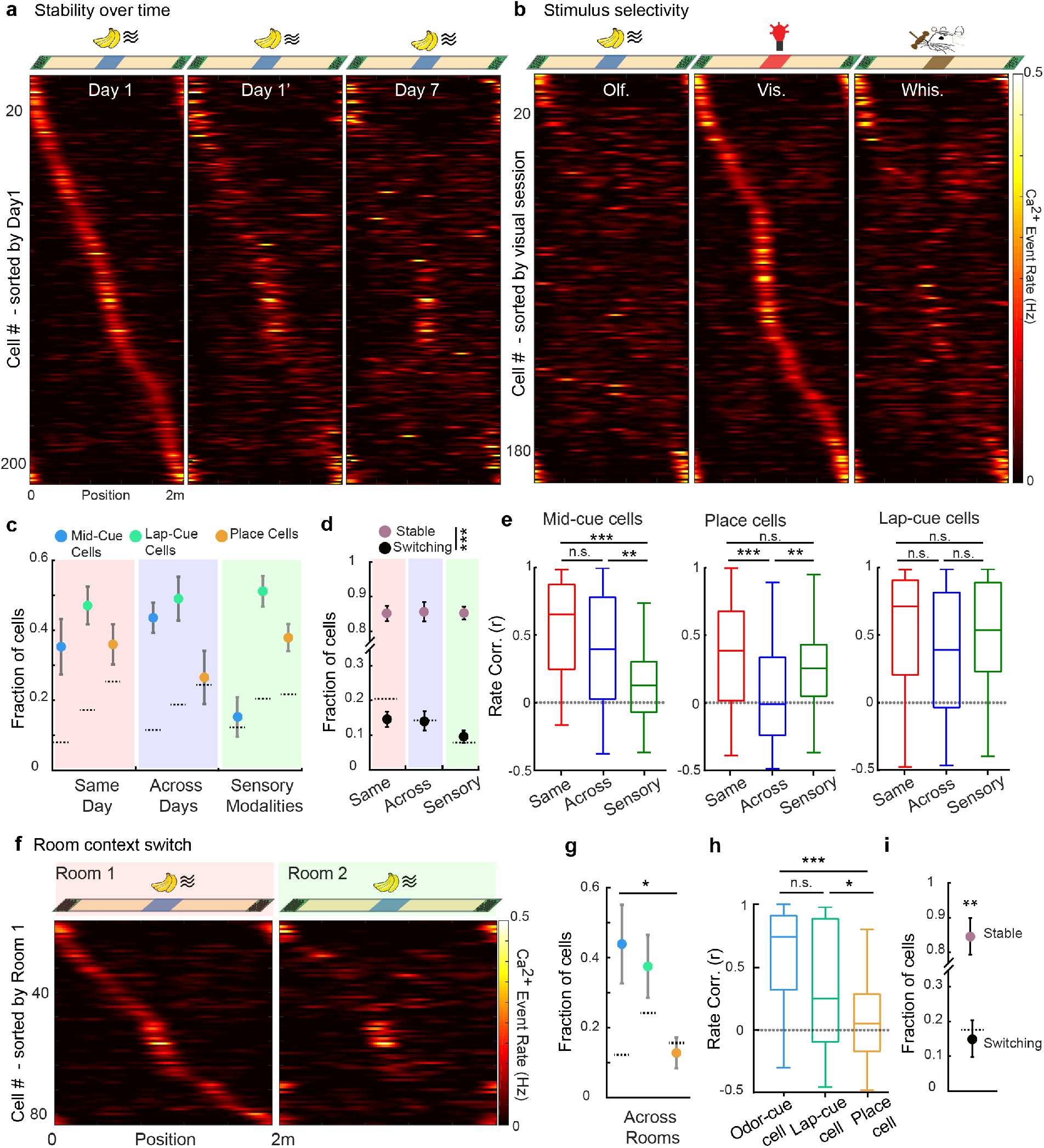
Cue cell responses are modality-specific, and more stable across days and room contexts than place cells. **a)** Spatial firing rates of individual DG granule cells matched between sessions within a day and after one week. Top: diagram of odor cue presentation during all sessions. Bottom: Spatial firing rates for spatially tuned neurons tracked during subsequent sessions on the same day or one week later, ordered according to the receptive field location of each cell during the first exposure (n=233 cells, 8 mice). **b)** Activity of granule cells in response to different sensory cues. Top: diagram of sessions with cues of different sensory modalities in the middle position but a similar lap cue. Bottom: Spatial rates for neurons tracked through consecutive sessions during exposures to different cues. Data are shown for neurons with significant tuning in the visual cue session and tracked in olfactory and whisker tactile cue sessions (n=196 cells, 6 mice). See also Figure S2a-d **c)** Fraction of cross-session registered cells that encoded the same behavioral variable, lap cue (green), middle cue (blue), or place (orange), within same day (left), across days (1wk, middle), and during exposures to different sensory modalities (right). Cells encoding the same variable in different sessions on the same day, χ^2^=1.81, p=0.4039, *P_LapCue-OdorCue_*=0.4221, *P_LapCue-Place_*=0.549, *P_OdorCue-Place_*=0.9713; cells encoding the same variable across days, χ^2^=14.36, p<0.001, *P_LapCue-OdorCue_*=0.1566, *P_LapCue-Place_*<0.0001, *P_OdorCue-Place_*=0.0313 (8 mice, 8 matched sessions, Day1-Day1’, Day1-Day7); cells encoding the same variable across different sensory modalities, χ^2^=19.78, p<0.00001, *P_LapCue-OdorCue_*<0.00001, *P_LapCue-Place_*=0.0766, *P_OdorCue-Place_*=0.0464 (6 mice, 12 matched sessions, Odor-Vis. and Vis.-Tact.); Kruskal-Wallis test, Dunn’s multiple comparisons test. Dashed lines represent 97.5^th^% of null distributions for each cell type. Error bars, mean ± SEM. **d)** Fractions of spatially tuned cells that stably encoded only one variable (cue or place, pink) or switched response types (cue to place or vice versa, black) within same day, across days and in response to different sensory cues: *P_Switch-Stable (Same Day)_*<0.001, *P_Switch-Stable (Across Days)_*<0.001, *P_Switch-Stable (Sensory Modalities)_*<0.001, Wilcoxon Rank Sum test. Dashed lines: 2.5^th^ and 97.5^th^% of null distributions. Error bars, mean ± SEM. **e)** Mean spatial firing rate correlations within the same day (red), different days (blue) and with different sensory cues (green) for mid-cue cells (left, χ^2^=27.88, p<0.0001, *P_Same Day-Across Days_*=0.2465, *P_Same Day-Sensory Modalities_*<0.0001, *P_Across Days-Sensory Modalities_*=0.0038, *N_Same Day_*=37, *N_Across Days_*=37, *N_Sensory Modalities_*=55); place cells (middle, χ^2^=18.79, p<0.0001, *P_Same Day-Across Days_*<0.0001, *P_Same Day-Sensory Modalities_*=0.7449, *P_Across Days-Sensory Modalities_*=0.0055, *N_Same Day_*=76, *N_Across Days_*=76, *N_Sensory Modalities_*=83), lap-cue cells (right, χ^2^=5.096, p=0.0782, *P_Same Day-Across_ Days*=0.0723, *P_Same Day-Sensory Modalities_*>0.9999, *P_Across Days-Sensory Modalities_*=0.7932, *N_Same Day_*=80, *N_Across Days_*=80, *N_Sensory Modalities_*=59). Kruskal-Wallis test, Dunn’s multiple comparisons test. Boxes, 25th to 75th percentiles; bars, median; whiskers, 99% range. \**P* < 0.05; \*\**P* < 0.01; \*\*\**P*< 0.001. **f)** Activity of DG granule cells in response to the same odor-cued track when recorded in different rooms. Top: diagram of treadmill track with an odor in the middle position and an invariant lap cue, performed in different rooms. Bottom: Spatial firing rates for tuned neurons tracked through consecutive sessions in different rooms (n=83 cells, 4 mice). See also Figure S5. **g)** Fraction of individual cells that encoded the same variable, odor cue (blue), lap cue (green) or place (orange) in different recording rooms. χ^2^=6.59, p=0.0372, *P_OdorCue-Place_*=0.0490, *P_LapCue-OdorCue_*=0.9557, *P_LapCue-Place_*=0.0972. Dashed lines: 2.5^th^ and 97.5^th^% of null distributions. Error bars, mean ± SEM. **h)** Comparison of spatial firing rate correlations in all tuned cells across rooms (χ^2^=15.85, p<0.001, *P_Odor-Lap cue_*=0.27, *P_Odor Cue-Place_* <0.001, *P_Lap Cue-Place_* =0.03, *N_Odor-Cue_*=25, *N_Lap-Cue_*=33, *N_Place_*=26). Kruskal-Wallis test, Dunn’s multiple comparisons test. Boxes, 25th to 75th percentiles; bars, median; whiskers, 99% range. \**P* < 0.05; \*\**P* < 0.01; \*\*\**P*< 0.001. **i)** Fractions of spatially tuned cells that stably encode only one variable (cue or place, pink) or switched response types (cue to place or vice versa, black) across different rooms (p<0.01, Wilcoxon Rank Sum test.) Dashed lines: 2.5^th^ and 97.5^th^% of null distributions. Error bars, mean ± SEM.

Between any two sessions, over days or with different cue modalities, the cells encoding the invariant lap cue were the largest fraction of cells that remained active and maintained their response type (e.g. cue type and/or place, green circles in Figure 3c). Odor cue cells fired reliably to the same olfactory cue over long periods of time (blue circles in red and blue shaded area, Figure 3c), but were largely unresponsive to cues of other modalities presented at the same position (green shaded area in Figure 3c, χ^2^=19.78, p<0.0001, Kruskal-Wallis test). Conversely, a lower percentage of place cells maintained their response type across days compared to cue cells recorded in the same sessions using the same cues (orange circles in blue shaded area, Figure 3c, χ^2^=14.36, p<0.001, Kruskal-Wallis test). Furthermore when we examined specifically whether cue cells become place cells between these sessions, or vice versa, we found that categorical cue and place representations remain extremely stable within the DG population (Figure 3d, p<0.001, Rank Sum test).

We further examined the degree of stability and specificity of cue and place cell tuning by cross-correlating spatial firing rates for individual registered cells over time and with respect to different sensory cues (Figure 3e). For this analysis, we computed the correlation of spatial firing rates for cells registered during different sessions within a day and one week later, or with a different sensory cue modality. In line with the stimulus selectivity analysis described above, we observed that middle location odor cue cells displayed significantly lower correlations in sessions with a different sensory cue, compared to separate sessions using the same olfactory cue on the same day or one week later (left, χ^2^=27.88, p<0.0001, Kruskal Wallis test). Thus, cells that responded to a cue of one modality were unlikely to respond to cues of other modalities, despite a similar spatial location of the cues. Correlations in the activity of individual place cells over 1 wk. were significantly lower than on the same day, and were lower than both cue cell populations, again indicating lower stability for place than cue representations (left, χ^2^=18.79, p<0.0001, Kruskal Wallis test). Lap cue cells did not display significant changes in their firing rate correlations between sessions on the same day, across days or with different sensory cues, and were therefore especially stable.

Responses of individual odor cue cells were also highly correlated in sessions recorded in different rooms on the same day (Figures 3f-h, S5h), indicating that distal ambient cues that differed between the rooms have a limited influence on the stability of sensory cue representation in DG granule cells. Despite the high stability of cue cells, place cells were significantly less consistent between rooms than cue cells in the same sessions (Figure 3g, h, p<0.001, Kruskall-Wallis test), and were similar to that seen in place cells over 1wk when measured in the same room (Figure 3e), suggesting that place cells are more context selective than cue cells. Furthermore, cross-registered place cells did not become cue cells in the other room, and vice versa (Figure 3i, p<0.01, Wilcoxon rank sum test). Taken together, these results suggest that sensory cues are represented by a stable subpopulation of neurons that is highly selective for specific cues while purely spatial representations are less stable and undergo progressive reorganization over time and in different global contexts.

### Spatial Modulation of DG Cue Responses

The juxtaposition of inputs from the lateral and medial entorhinal cortex onto the dendrites of individual granule cells has been hypothesized to underlie a conjunctive code for sensory cues and their spatial location^3,4^. We therefore examined the influence of spatial location on cue responses (“spatial modulation”) by tracking odor cue cells through multiple sessions with different cue-location pairing, either with an intermittently shifted cue as in previous experiments or in separate sessions with the same cue at random locations each lap (Figure 4a, b). On average, cue-triggered Ca^2+^ response amplitudes for individual cue cells (a proxy for neuron action potential burst firing rate) were smaller when cues were presented at the infrequent “shift” location or at random locations, when compared to their responses at the more frequent middle location (Figure 4c, d, p<0.0001, Friedman test, n=101, 5 mice). Thus while cue cells tend to respond to the same sensory cue regardless of location, the strongest responses occur when a cue is presented at the same place repeatedly.

**Figure 4:**
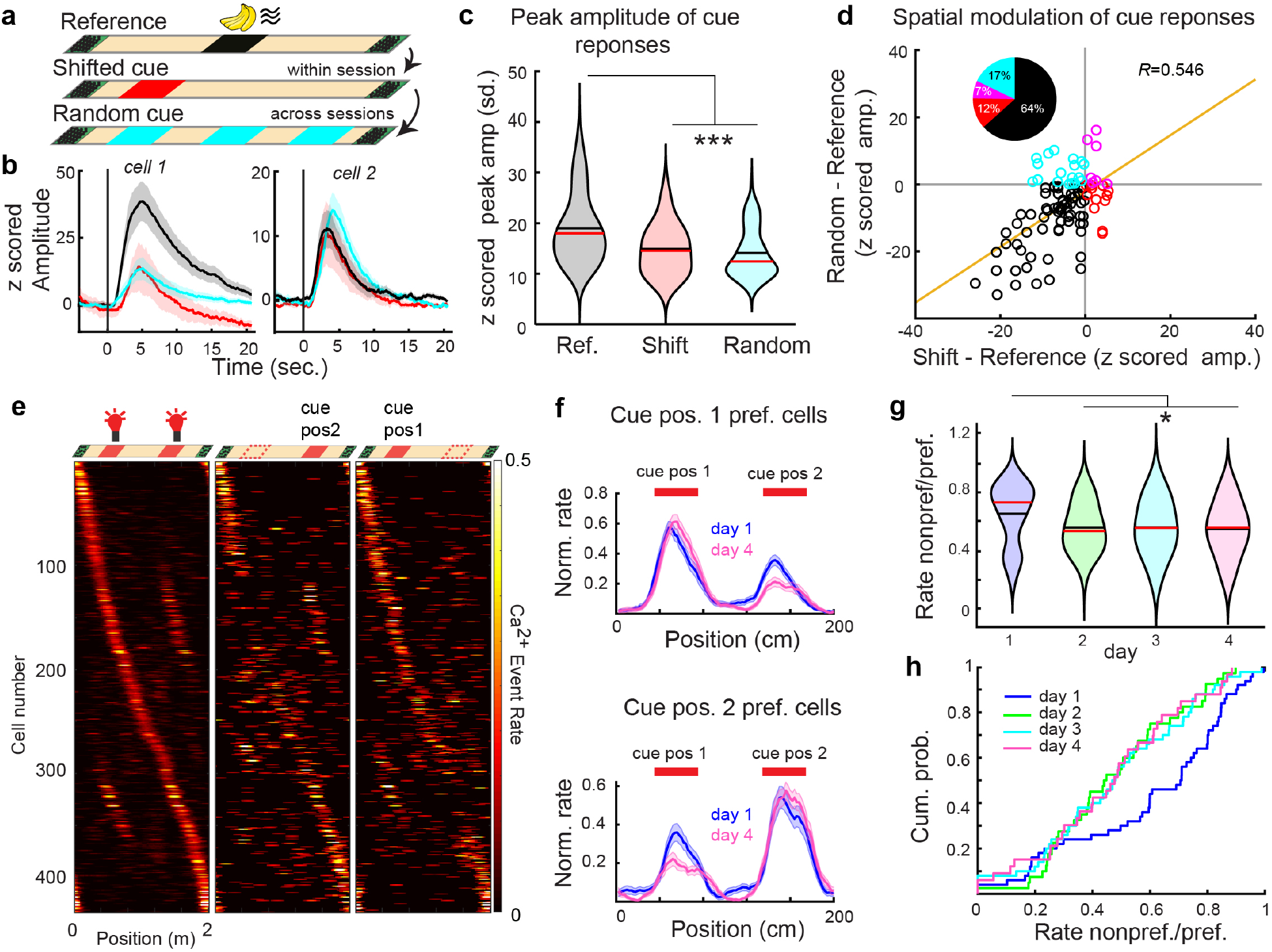
Sensory cue responses are spatially modulated. **a)** Experimental setup to examine effects of cue-location pairing on cue responses. Odor cue response amplitudes are measured for the more frequent middle cue location (reference, black), an intermittently shifted cue location in the same sessions (shifted cue, red), or with random administration of the same cue at one of three positions on each lap in a subsequent session (random cue, cyan). **b)** Example cue-triggered average Ca^2+^ transients from two example cue cells on normal (black), cue-shifted (red) and random-cue (cyan) laps/sessions. **c)** Peak amplitudes of Ca^2+^ transients from individual odor cue cells in normal, shift and random cue presentation conditions; χ^2^=49.36, p<0.001, *P_Normal-Shift_*<0.0001, *P_Normal-Random_*<0.0001, *P_Shift-Random_*>0.9999, Friedman test and Dunn’s multiple comparisons tests (n=101 cells, 5 mice). Red lines in violin plots show median, black lines show mean. **d)** Difference in the magnitude of cue responses in normal laps from random-cue and cue-shift laps. Yellow line, linear regression (R^2^= 0.546, p=3.5×10^−9^). Pie chart shows the percentage of neurons with higher average responses in normal (black, 64%), cue-shift (red, 12%), random cue presentation conditions (cyan, 17%), and both shifted and random laps (purple, 7%). **e)** Dual cue location task: (Top) An LED visual cue is given consistently at two positions on the track. Arrangement of cues for normal laps (left, 80% of laps), laps where first cue is omitted (middle, 10%), and laps where the second cue is omitted (right, 10%). (Bottom) Spatial firing rates of 433 significantly tuned cells on first day of task (N=3 mice). **f)** Average spatial firing rates for significant cue cells, for cells preferring cue position 1 (top) or cue position 2 (bottom) on first day of task (blue, n=25 pos1, n=25 pos2) and 4th day of task (pink, n=13 pos1, n=20 pos2 cells). **g)** Spatial modulation index (rate non-preferred cue/preferred cue) for all cue cells on days 1-4. (p=0.0125, day 1 vs. days 2, 3, or 4, n= 50/40/50/33, Wilcoxon rank sum test). Note that lower values mean higher cue spatial selectivity. **h)** Empirical cumulative probability distribution of spatial modulation index for all cue cells on days 1-4. See also Figure S6a-d.

To further examine the spatial modulation of cue responses, we performed a set of experiments where a visual cue was delivered at multiple locations on the track to measure selectivity of granule cell responses to cues presented consistently at distinct locations (“dual location cue”, Figure 4e), similar to recent work by other groups using virtual reality environments^41,42^. As in our previous experiments with a single cue, each cue was omitted on a subset of laps to isolate cue cells as opposed to place cells present in the vicinity of cues (Figures S6a, b). To measure the extent of preference for one cue location versus the other, we calculated the spatial modulation index for each cue cell as the ratio of the firing rates in the non-preferred cue location versus the preferred cue location. Cue cells in the first session exhibited a range of spatial modulation indices, but over the population their event rates were significantly modulated by the cue location (p<0.00001, Wilcoxon signed-rank test on day 1 spatial modulation index, 53% of individual neurons significantly spatially modulated by location shuffle, Figure 4f). Spatial modulation also increased over several days of dual cue presentations (p=0.0125 Wilcoxon rank sum test, day 1 vs 2, 3, or 4), with 81% of cue cells exhibiting significant spatial modulation by day 4 (Figures 4-h, S6c-d). This suggests that dentate cue cells are acutely modulated by the spatial location of two identical cues but increase their preference for a single location over days.

### Effects of Cue Manipulation on Spatial Encoding in the DG

While the most robust activity in the DG was found within the cue cell population, the existence of the smaller population of place cells in uncued locations suggests that spatial activity in these cells is referenced to one of the two cues on the otherwise featureless treadmill track in order to encode a unique location. We therefore examined the effect of manipulations of the variable cue on the subsequent spatial encoding of place cells, in order to judge the degree to which this cue acts as a landmark. For example, if place cells were acutely referenced to the nearest cue we would expect the place fields of all cells following the variable middle cue to shift on cue shifted laps. To examine the relationship between spatial firing patterns in normal cue laps to those in cue shifted laps, we first calculated population vector (PV) correlations of firing rates across all spatially tuned cells (cue and place cells) on each lap (Figure 5a). While the correlation was higher in the vicinity of the cues between normal laps, we observed a dramatic decrease in PV correlation between normal and cue shifted laps which was confined to the area immediately around the cue itself. However, this change in PV correlation did not extend much beyond the cue location, consistent with a limited effect of cue shifting on subsequent place cell activity.

**Figure 5:**
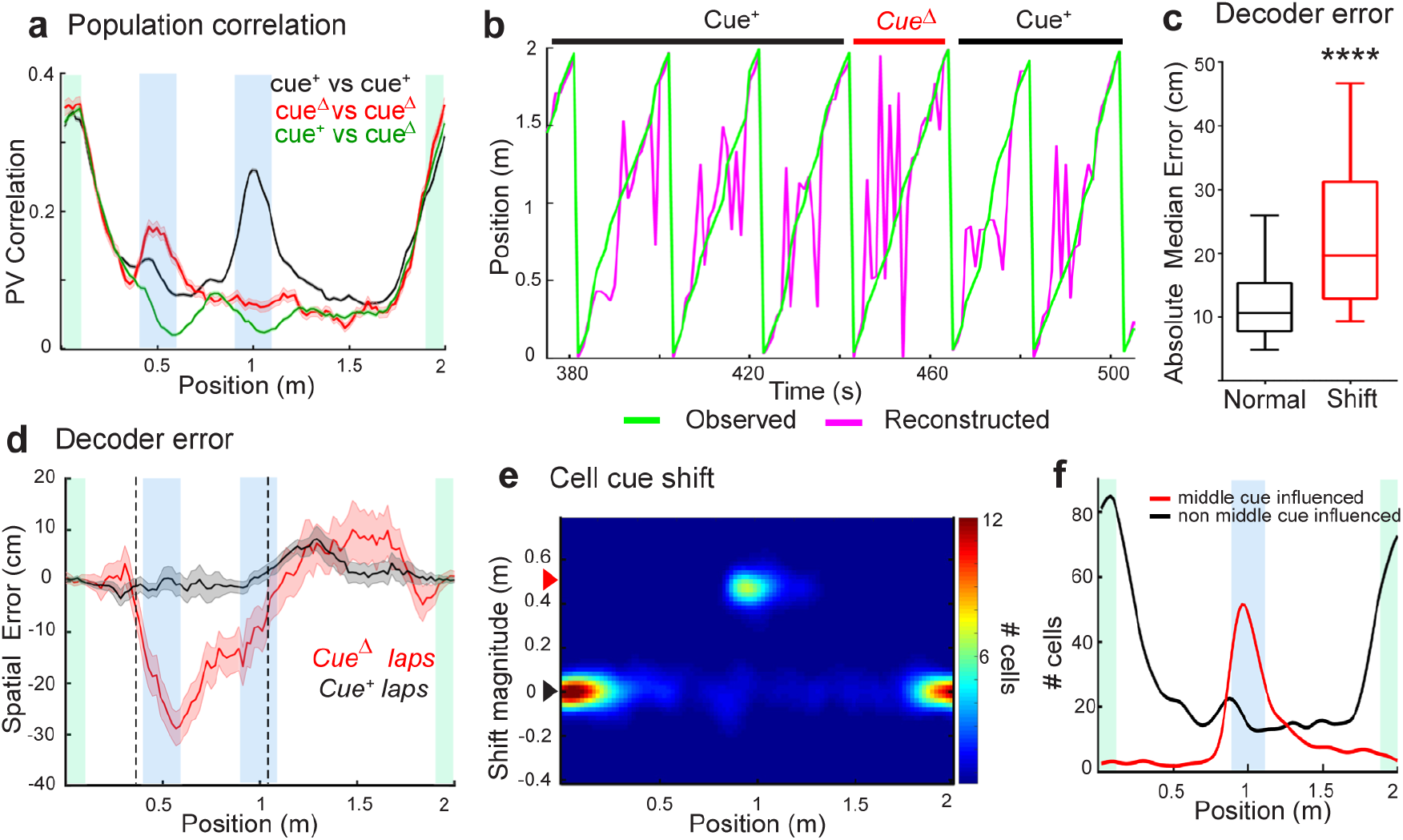
Effects of cue manipulation on spatial encoding. **a)** Population vector correlations for all spatially tuned cells during normal middle cue laps and shifted cue laps. Pair-wise correlations of normal middle cue laps are shown in black, cue shifted laps are shown in red and between normal and shifted laps are shown in green. Large differences in pair-wise correlations are largely confined to locations of the middle/shift and lap cue that are shown blue and green shaded areas, respectively. **b)** Bayesian decoding: example prediction of 4 normal and 1 cue shifted laps for a representative session, based upon activity from all spatially tuned granule cells. The magenta line shows predicted position at each time bin, while the green line shows the observed position of the animal. **c)** Average prediction error for normal and cue shifted laps calculated as the absolute median distance between the decoded value in each time bin and the actual value of the position (p<0.0001, Wilcoxon signed-rank test, n=66 sessions). Boxes, 25th to 75th percentiles; bars, median; whiskers, 99% range. **d)** Spatial decoding error for each treadmill position in normal (black) and cue shifted laps (red). Shaded error region represents SEM of the difference of the predicted position from the animal’s actual position on the treadmill by session. Dotted lines indicates point of statistical equality (*p*>0.05, Wilcoxon Rank Sum test) between normal and cue shifted lap decoding. **e)** Single-neuron receptive field shifts: Distribution of firing location shift on shifted cue laps for all spatially tuned cells, plotted based upon tuning location in normal middle cue laps (n=66 sessions). Red arrowhead indicates actual cue shift distance, and thus marks cells that are directly influenced by the cue. Black arrowhead indicates cells with zero shift, and thus not influenced by the odor cue. **f)** Mean number of cells not influenced by the odor cue (shift magnitude<0.05m, “non middle cue influenced”) and cells shifting precisely along with the cue (shift magnitude 0.5m+/−0.1m, “middle cue influenced”) at each track position (corresponding to cells in regions of black and red arrowheads in “e.”, respectively).

To further evaluate the manner in which cue associated activity contributes to spatial encoding, a spatial Bayesian decoder was constructed from the firing rate vectors of all spatially tuned cells (4,091 cells from 66 sessions)^27^. In order to establish a non-biased estimate of the position during treadmill running, the decoding was performed using a 5 fold cross validation approach in which 1/5^th^ of decoded laps were held out from the training set. Post-reconstruction, we divided the data according to the lap types (Figure 5b). The decoder accuracy was higher in normal (median, 10.6 cm) compared to both cue shift (median, 19.6 cm) and cue omitted laps (median, 18.4 cm, p<0.0001, Wilcoxon rank sum test, Figure 5c, data not shown for omit laps), indicating that cue manipulations affect the accuracy of spatial coding by the DG population. However while the cue shift strongly perturbed spatial decoding near the shifted cue, estimating the animal’s position to be farther along than the real position, the decoder error on shift laps soon converged to that of normal laps, well before the subsequent lap cue (Figure 5d, dotted line). Together these results suggest that as a population, the spatial encoding of granule cells is affected by sensory cues within a localized area near the cues, rather than persistently altering their estimation of the animal’s position like a landmark.

Population vector correlations and Bayesian decoding are average measures of spatial representation in the granule cell population as a whole, which might obscure differences in how the activity of individual cells is referenced to local or distant cues. We therefore examined responses of individual granule cells by measuring cross-correlation offsets between spatial firing in normal and cue-shifted laps for each cell. In doing so we found two distinct classes of single-cell shift responses: some cells shifted their firing precisely along with the odor cue shift distance (0.5m), while others were consistently referenced to the stable lap boundary cue and did not shift their response location during the odor cue location shift (Figure 5e, f). The majority of cells that shifted their firing in reference to the new cue location were active immediately in response to the middle cue (“middle cue-influenced”), while a small number were persistently affected by the cue shift. In contrast, a relatively constant number of place cells along the track maintained their normal firing location with reference to the distant lap boundary cue despite the shifted local cue (“non-middle cue-influenced”). Notably, there was also a small enrichment of place cells immediately prior to the cue but unaffected by the cue shift (Figure 5f), indicating a spatial response predictive of the normal cue location. The presence of two distinct groups of cells that fire in reference to the two cues on the treadmill track (the variable middle cue and stable lap cue) indicates that the DG population maintains independent spatial reference frames based upon distinct cues within an environment. This code however appears to differ depending upon the spatial stability of the cues: the effects of spatially variables cues are limited to the cue cell population while place cells are referenced to global landmarks^43^.

### Cue-related Inhibition of DG Responses

Spatial receptive fields of granule cells are hypothesized to form within a competitive network in which lateral inhibition mediated by GABAergic interneurons enforces sparse encoding that may aid pattern separation^17,44^. The robust and highly selective cue responses of granule cells in our behavioral paradigm allowed us to examine patterns of local-circuit inhibition within the DG network. First, we sought to analyze the effect of cue responses on spontaneous firing observed within the spatially tuned granule cell population. By comparing spontaneous firing rates at the middle cue location outside of the primary spatial receptive field in normal laps (“out-of-field”) with laps where the cue is omitted, we identified a significant reduction in background firing of the spatially tuned granule cell population during the cue presentation (Figure 6a, p=0.0004, Wilcoxon signed-rank test). This suppression was absent on cue-omitted laps and generally co-varied with the mean amplitude of cue-related excitation (within-field) on a session-by-session basis (Figure 6b, R=0.22, p<0.0001, Wilcoxon signed-rank test). Furthermore, the timing of the peak of this inhibition was delayed with respect to the excitatory cue response in these sessions (Figure S6e, f). These data show that sensory cue input is able to suppress ‘noisy’ spontaneous firing in the granule cell population, potentially through lateral inhibition.

**Figure 6:**
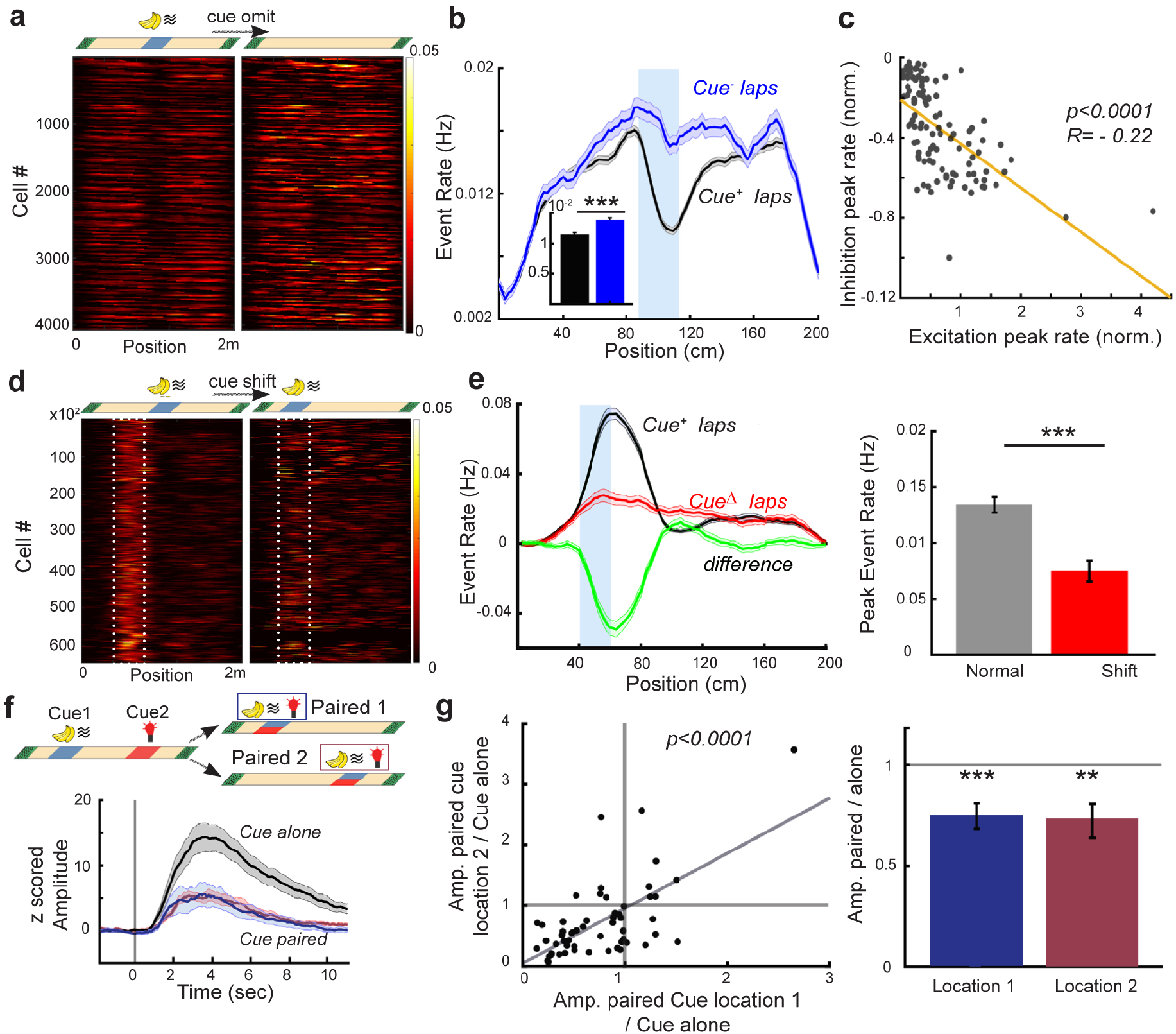
Cue presentation leads to suppression of diverse types of DG responses. **a)** Cue-related suppression of spontaneous, out-of-field firing of DG neurons: Spatial firing rates outside of the spatial receptive field center-of-mass (+/−10cm) for significantly tuned cells during normal and cue omitted laps (n=4,091 cells from 66 sessions). **b)** Average out-of-field, spontaneous firing rates from all spatially tuned cells in the presence of the middle cue (black) compared to laps in which the cue is omitted (blue), within the same session. Blue shaded area shows the cue delivery position. (Inset) Mean ± SEM of firing rate within cue region, p<0.001, Signed-rank test, n=4,091 cells. **c)** Comparison of the magnitude of cue-related excitation and inhibition by session: Average in-field spatial firing rates (i.e. cue-related excitation), compared with out-of-field, spontaneous firing rates (i.e. cue-related inhibition) at the middle cue location for each session (90-120cm), adjusted for pre-cue firing rate (66 sessions). Yellow, linear regression: p<0.0001, R^2^=−0.22. **d)** Cue-related suppression of place cell firing: Spatial firing rates of place cells with firing fields at 50-80cm during normal laps (left) and laps in which the cue is shifted to the same 50cm location (right). **e)** (Left) Average spatial firing rate for the above 50-80cm receptive field place cells in normal laps where the cue is not presented at this location (Cue+, black), compared with laps in which the cue is shifted to this location (CueΔ, red; diff. in green). (Right), Average peak firing rate of the same cells in “d.” and “e.”, paired t-test, p<0.001, 645 cells. **f)** Cue-related suppression of cells responding to other distinct cues: Top: Diagram of intermittent cue pairing experiment. Cues of two different modalities were presented at different locations, interspersed with paired presentation at one of these locations on intermittent laps. Bottom: Example cue-triggered averaged Ca^2+^ transients for a cue cell strongly active when the cue is presented alone (black) but with reduced responses when paired with a different cue, regardless of location (blue, pair location #1, purple, location #2). **g)** (Left) Amplitudes of paired cue responses for individual cue cells at the two pairing locations with respect to the response to the cue alone. Gray line represents the diagonal, p<0.0001, R^2^=0.91, n=56 cells, 3 mice. (Right) Average relative response amplitude at the two pairing locations (Location 1: 0.75± 0.061, p1 = 1.45×10^−4^; Location 2: 0.73±0.086, p2= 0.003, Wilcoxon Signed Rank Sum test, bar plots are mean ± SEM).

In addition to suppressing spontaneous firing activity in the DG network, we sought to determine the effect of cue responses on spatially tuned activity in the place cell population and in cue cells responding to other differing cues. By examining cells with receptive fields normally at the intermittent shift location, we found that place cell firing rates were strongly suppressed on shift laps when a cue was presented at this location compared to normal laps when a cue was not present here (Figure 6d, e). Furthermore when two distinct cues were normally presented at different locations, intermittent shifting of the position of one cue to overlap with the other cue typically had the effect of suppressing the responses in both cue populations (Figure 6f, g), indicating mutual inhibition among granule cells encoding different cues. Thus in addition to the inhibition of spontaneous firing in the granule cell population, cue responses lead to overall inhibition of place cell responses and responses to competing cues as well. Taken together, our results suggest that cue-related activity in granule cells and the resulting suppression of responses in neighboring cells via lateral inhibition may organize the dentate gyrus into a competitive network that utilizes the strongest and perhaps the most informative parameters, such as stable landmark cues, in order to accurately establish an animal’s location in space.

## Discussion

By imaging calcium activity in large populations of dentate granule cells during head-fixed spatial behaviors on a treadmill, we have shown that a major population of task-selective neurons is highly sensitive to specific sensory cues (non-spatial, or ‘what’ information) rather than to discrete locations (spatial, or ‘where’ information). The ability to dynamically manipulate a sensory cue and its association with locations on the treadmill track allowed us to isolate the population of cue cells, in addition to the complementary population of place cells recorded in the same sessions.

Properties of cells within these groups also differed: sensory cue responses were highly-tuned and remarkably robust across contexts and over time-significantly more so than the responses of the canonical place cell population (Figures 1–3). And while cue cells did not have spatially tuned responses in prior sessions on the same treadmill track without the odor cue (Figure S4), or in the first several cue presentations, they emerged rapidly during the first session of exposures to the cue while place cell responses emerged more slowly (Figure 1i). These findings suggest that dentate cue cells stably represent both sensory cue and other non-spatial features in an environment (e.g. reward, Figure S2e-h), while place cells constitute a more dynamic population that gradually adapts to current conditions, for example, by integrating between stable cues in order to provide an accurate estimate of the animal’s location when no cues are present^45^.

### Heterogeneous properties of Granule Cells

The robust and consistent nature of GC cue responses suggests that cue-selective populations may have been similarly present in previous experiments recording DG activity *in vivo*, however the lack of precise stimulus control made it impossible to distinguish cue-responsive versus place-responsive components^5,7,36,46^. Indeed, spatial receptive fields of individual granule cells measured in different environments with similar sensory cues display mixed levels of changes in their firing rates, with a subset of granule cells completely remapping their firing fields while others show more stable dynamics^5,44,46–48^. Based upon our results, these subgroups may correspond to place cells and cue cells, respectively. A major impact of sensory cues on DG firing largely independent of spatial context might also explain the relatively lower context selectivity observed in the DG compared with other hippocampal subfields, at least when measured in different contexts that contain the same or similar sensory cues^5,44,46–48^. To support this notion, we found that DG cue cells were largely stable when recorded over long periods of time, with different cue positions, locomotion speeds, or in different room contexts, while place cells were not (Figures 2–4). Previous work has demonstrated that the two major long range inputs to the dentate gyrus, the lateral and medial entorhinal cortex (LEC and MEC), are involved with processing functionally distinct information^10,11^. This raises the possibility that our “cue cells” may be driven primarily by sensory inputs from the LEC, while “place cells” are driven by self-motion information relayed from the MEC, for example in the form of grid cell responses. The segregation of these properties within the overall granule cell population suggests that these streams of information remain largely separated at the level of the DG^49^, similar to the functional heterogeneity observed in other hippocampal subfields^50,51^. In these regions such heterogeneity among principal neurons has been hypothesized to be critical for efficient encoding of spatial representations within a local population^49–52^. Heterogeneity has also been observed in immediate early gene expression profiles within DG neurons encoding individual memory engrams^53^ which are differentially targeted by entorhinal afferent pathways^9,53–55^. These findings are in line with our study demonstrating that sensory and spatial information remain partially separate at the level of the DG, and suggest that multiple channels of spatial and non-spatial information contribute distinctly to local and down-stream computations. This separation is also consistent with current ideas that cue-based and path-integration based navigation are complementary, rather than integrated, in order to produce place-specific firing both in locations where landmark cues are present as well as in between cues^56^.

Motorizing the treadmill, while different from previous studies^5,7,23,44^; does not affect our central claim of the strong representation of sensory cue information in the DG (Figures 2 and S1). Additionally, running on motorized treadmills does not influence the differences in the stability and tuning properties between the cue and place encoding populations (Figures 1 and S1). Yet, we note that these differences obtained during head-fixed navigation on a relatively cue impoverished track at a constant speed may be larger than what would be observed during self-guided exploration of cue-rich, multisensory real-world environments. Nevertheless, our ability to track the same neurons across different locomotion conditions provides additional evidence for the existence of a separate and non-overlapping population of cue cells in the DG compared to the ‘classical’ place cells reported in previous studies^5,7,23,44,47,48^. Indeed, we have found repeatedly that cue cells displayed striking stability under different imaging conditions (locomotive behaviors, across days, different rooms), while the spatial fields of place cells remapped in the same set of conditions, suggesting that place cells integrate position and contextual information to a higher degree than cue cells. Thus, we reason that the classification of individual cue cells and place cells in the DG would likely remain consistent in more complex environments such as real-world or visuospatial virtual environments. Future studies investigating whether cue and place encoding populations receive different inputs from LEC and MEC, as well as the molecular basis of the heterogeneity among granule cell populations, will allow for a more detailed inquiry into the functional diversity in DG granule cells. Such studies may also allow us to selectively target cue and place cells in order to manipulate their activity and determine their roles during discrimination behaviors.

### Functional significance of dentate gyrus cue cell properties

But how do these results inform our ideas about how the DG participates in pattern separation and spatial map formation? First, we found that although individual granule cells respond to the same cues when presented in different locations, the strength of this activity was spatially modulated (Figures 4, S6). Differential encoding of cues based upon their spatial location may contribute to pattern separation by allowing discrimination between similar cues present in different places within an environment, or in distinct contexts, over the population. We observed that spatially modulated cue responses also increased over days when a single cue was repeatedly presented at multiple locations (Figure 4h), indicating plasticity in location specificity and potentially progressive enhancement of spatial pattern separation. Notably, responses were also on average larger for odor cues presented at a single location, as opposed to shifted or random locations (Figure 4b-d). This suggests that cue responses are more robust for stimuli paired repeatedly with one particular location, which may indicate dendritic integration of sensory and spatial information. Conjunctive encoding of sensory cues and their locations within an environment could play a role in establishing landmarks for spatial navigation. In support of this idea, we found that most place cells were referenced to the stable lap cue rather than the variable middle cue, indicating that this cue is preferentially utilized as a landmark (Figure 5).

Our study also shows that cue responses lead to potent inhibition of three distinct types of granule cell activity: spontaneous “noisy” spiking activity, the spatial firing of place cells, and the activity of cells responsive to other overlapping sensory cues (Figure 6). This supports the hypothesis of a competitive network in the DG enforced by strong lateral inhibition, which has been suggested to contribute to pattern separation. In addition, suppression of weaker place cell responses by cue-related activity may indicate that the network is organized to utilize the strongest and perhaps the most informative parameters, such as landmark cues, in order to establish an animal’s location in space.

Taken together, rate modulation of granule cell cue responses resulting from spatial location or association with other cues is itself a form of “rate remapping”, a feature frequently attributed to the dentate gyrus in pattern separation^3,17,19,46^. Thus spatial location (Figures 4c,d, 5e,f), as well as the juxtaposition of multiple cues (Figure 4e,f) and their associated patterns of inhibition (Figure 6), may create a rate modulated landscape of granule cell activity specific to the current context. The overall pattern of contextually modulated cue responses in the DG may form the basis for recruitment of distinct populations of neurons, or “global remapping”, observed in area CA3^46^. Slower emergence of the place cell population, as well as lower stability over sessions and in different room contexts, may also point to a role for these neurons in context selective encoding of position based upon integration between current landmarks (Figures 1–3).

While previous work has shown that CA regions also have cue responses^31–35^, all of these areas receive direct (CA3) or indirect input (CA1) from the DG as well as entorhinal cortex, so it remains unclear whether these properties arise as a result of direct inputs from entorhinal regions, or through the hippocampal trisynaptic circuit. Thus, the DG is the best place to look specifically at how the presumed different channels of information from the lateral and medial entorhinal cortex are initially integrated by the hippocampus during spatial map formation. Our work suggests that the DG encodes representations of sensory cues in the environment that interact weakly with spatial representations, potentially through lateral inhibition. The rate-modulated code of DG sensory cue representations could be refined in subsequent hippocampal areas such as the CA3 recurrent attractor network which may resolve conflicting cue and path-integration based information to encode an integrated spatial map of things and places, important both for spatial navigation as well as episodic memory. Future studies examining how these specific features manifest in subsequent stages of the hippocampal circuit, will shed light on the progressive processing and integration of sensory cue and spatial information in hippocampal computation.

The properties of cue and place cells in the dentate gyrus have the hallmark of features one might expect from the first stage of spatial map formation in the hippocampus: mostly distinct but slightly mixed encoding of sensory cues, or “what”, and place, or “where”. Such complementary channels of information may be further refined by downstream hippocampal areas into an integrated spatial map that encodes relationships between important features such as spatial cues and goals, which can be used to organize adaptive behavior. Uncovering the functional heterogeneity of cue and place cells in the dentate gyrus helps us to better understand the basic neural mechanisms underlying our ability to navigate in complex environments by utilizing both landmarks and self-motion information to guide our movements through space.

## Acknowledgements

We would like to thank Jack Berry and other members of the Hen lab for critical insight throughout this project. This work was funded by NIMH T32 MH015144, S10 OD018464, NIH K99 MH12226 (S.N.T.), NYSTEM-C029157 (G.O., G.F.T., C.O.L, R.H.), Revson Senior Fellowship in Biomedical Science (A.D.G.), NIMH R21MH122965,NARSAD Young Investigator Award (G.F.T.), NIMH R01 AG043688 (R.H.), MH068542 (R.H.), NIMH R01 MH100631 (A.L.), NINDS R01NS094668 (A.L.), and NINDS U19NS104590 (A.L.).

## Author Contributions

Conceptualization, C.O.L., S.N.T., and R.H.; Methodology, C.O.L., S.N.T., A.S.; Software, C.O.L., S.N.T., A.D.G., J.B.; Formal Analysis, S.N.T., C.O.L.; Investigation, C.O.L., S.N.T.; Resources, C.O.L., A.D.G., G.F.T., J.B., A.L., G.O.; Writing, Review and Editing, S.N.T., C.O.L., R.H., A.D.G., A.L.; Supervision, R.H and A.L.

## Experimental Procedures

### Mice

All procedures were conducted in accordance with the U.S. NIH Guide for the Care and Use of Laboratory Animals and the Institutional Animal Care and Use Committees of New York State Psychiatric Institute and Columbia University. Adult male C57BL/6J mice were supplied by Jackson Laboratory and Dock10^Cre^ mice were a gift from Susumu Tonegawa (Massachusetts Institute of Technology). Mice were housed in a vivarium grouped 2-4 mice/cage enriched with running wheels, maintained on a 12-hour light cycle and used at 8-10 weeks of age. Experiments were conducted during the light portion of the cycle. Food and water were available *ad libitum* until the beginning of the experiment, when they were placed under controlled water supply and maintained at >90% of their pre-deprivation weight over the course of imaging experiments. In total, imaging data from 18 mice were used in this study.

### Surgery

Dentate gyrus virus injection and imaging window implantation surgeries were performed as described previously (Danielson 2016, 2017). For all surgical procedures, mice were anesthetized with 1.5% isoflurane at an oxygen flow rate of 1 L/min, and head-fixed in a stereotactic frame (Kopf Instruments, Tujunga, CA). Eyes were lubricated with an ophthalmic ointment, and body temperature maintained at 37°C with a warm water recirculator (Stryker, Kalamazoo, MI). The fur was shaved and incision site sterilized prior to beginning surgical procedures, and subcutaneous saline and carpofen were provided peri-operatively and for 3 days post-operatively to prevent dehydration and for analgesia. Mice were unilaterally injected with recombinant adeno-associated virus (rAVV) carrying the GCaMP6s transgene (pAAV.Syn.GCaMP6s.WPRE.SV40) purchased from Addgene (viral prep #100843-AAV1) with titer of 1-5×10^12^ in dorsal dentate gyrus using a Nanoject syringe (Drummond Scientific, Broomall, PA). Injection coordinates were −1.5 mm AP, −2 mm ML, and −1.85, −1.7, −1.55 mm DV relative to the cortical surface. 30 nL of diluted virus was injected at each DV location in 10 nL increments. Mice were allowed to recover for 3 days and then were unilaterally implanted with an imaging window and stainless steel head-post for head fixation. Imaging windows were constructed by adhering 2 mm diameter, 2.3 mm long stainless steel hypodermic tubing (Ziggy’s Tubes and Wires Inc, Pleasant Hill, TN) to 2 mm diameter glass coverslips (Potomac Photonics, Halethorpe, MD). A 2 mm diameter craniotomy was made centered on the previous injection site with a taper pointed-drill (Henry Schein Inc, 9004367) and dura was removed with micro curette (FST, 10080-05). The overlying cortex was gently aspirated to reveal capsular fibers with continuous irrigation with ice cold aCSF solution and bleeding was controlled with a collagen gel sponge (Avitene). Under minimal bleeding, a 30g blunt syringe was used to gently aspirate capsular and CA1 alveus fibers with white appearance and CA1 pyramidale and moleculare with pink appearance until vasculature of the hippocampal fissure became visible (under bright light with low bleeding). The cannula, attached to the stereotactic handle, was then gently lowered into the craniotomy and affixed to the skull using dental cement (Unifast Trad powder and LC light cured acrylic UV, Henry Schein).

### Behavioral training and apparatus

After a minimum of 1 week recovery period, mice underwent a water restriction scheme (1ml per day) and trained to run on treadmill while head-restrained. The training period typically lasted a week (2 training sessions/day, 15 min each) until the mice were able to run for at least 1 lap/ minute and seek reward from 3 randomly placed reward zones by licking the water delivery port. We then initiated the motorized belt adjusted to the natural velocity of each mouse and proceeded training for 1-2 more days. We discarded mice that did not perform sufficiently to receive getting all of their daily water supply during treadmill training, and were not motivated to move on the treadmill. A separate cohort of mice ran at will on the treadmill (Figures S1 and 2). During the training period mice ran on a cue-less ‘burlap’ belt and progressed to a different belt containing cues of different modalities as described below. For room switch experiments (Fig. 3f-h), mice were transferred to a separate room with similar imaging equipment on another floor of the building in their home cage. Mice were imaged for two consecutive sessions/day, each 15 minutes long, for the duration of 7-10 days. Single odor, light, and whisker texture responses described in Figures 1 and 3 and S2-5 are from the first time mice were exposed to these sensory stimuli. The subsequent exposures to these sensory cues, they were either in different locations on the belt, preceded or followed by additional cues (Figures 4, 6), reducing the effects of learning the location of these cues. Additionally, in all experiments, the treadmill belt material was changed between sessions to reduce the chances of urine contamination which might act as an additional olfactory cue.

The behavioral apparatus consisted of 2m long, 3” wide fabric belt stretched between 6” diam. laser-cut plastic wheels, mounted on an aluminum frame (8020.net). Spatial triggering of task events was performed by custom software via serial communication with a microcontroller (Arduino DUE) and an associated printed circuit board (OpenMaze OM4 PCB, https://www.openmaze.org) on the treadmill. The axle of the treadmill wheel was attached to a quadrature rotary encoder (US Digital part #: MA3-A10-125-B) connected to a custom quadrature decoder board and Arduino Nano (courtesy of Wen Li). Angular displacement was converted into a virtual linear distance based on the circumference of the treadmill. The errors were corrected via a registration anchor marked by radio-frequency identification (RFID) buttons (SparkFun Electronics) at the lap boundary of the belt and was detected when it passed over an RFID reader (ID-12LA, SparkFun) affixed underneath the mouse. A 12V DC gear motor was attached to the axle of the treadmill connected to a separate Arduino/OpenMaze shield using pulse-width modulation to adjust the rotation speed. A water reservoir connected to a water delivery port consisting of a small gavage needle (Cadence Science) was placed within reach of the mouse’s tongue. A capacitance touch sensor (Sparkfun MPR121) was attached to the water port to measure licking and the sensor was connected to the Arduino/OM4 PCB. Small 2-3ml drops of water were delivered by the brief opening a solenoid valve (Parker Hannefin) connected to the water port. Rewards were triggered at random locations each lap when mice entered a 10cm long reward zone on the track and were available until mice exited the reward zone or 3 sec had elapsed. Olfactory stimuli consisted of undiluted isoamyl acetate (IAA, Sigma W205532) which was added to syringe filters (Whatman #6888-2527) and delivered by opening a solenoid valve (SMC) connected to a flow controller delivering constant airflow of compressed medical grade air for 1s (~3psi). Visual and tactile stimulation consisted of a red LED contralateral to the imaged region, or a 1” square piece of sand paper brushed by the contralateral whiskers using a stepper motor, at approximately the speed of the treadmill belt. Custom written Be-Mate algorithm implemented in Java was used for recording mice’s licking, its position on the belt, and cue delivery. Mice were monitored using an IR camera (PS3eye) and illuminated using an IR LED array.

To isolate cue-selective responses among the granule cell population, normal cue laps in which the olfactory, visual, or tactile cue was presented in the middle of the treadmill track (90-110cm) were interspersed with occasional laps (10-20% of laps) in which the same cue was omitted (“omit” laps), or shifted forward ¼ of the track (“shift” laps). For a subset of sessions, the olfactory cue was presented at one of 5 locations along the track randomly each lap, in order to examine the effect of spatial pairing of the cue.

### In vivo two-photon imaging

Imaging was conducted using a microscope setup which consists of 8kHz resonant galvanometer (*Bruker*) mounted to a mirror-based multi-photon microscopy system (*Prairie Technologies*) and an ultra-fast pulsed laser beam (920-nm wavelength; *Chameleon Ultra II, Coherent*, 20–40-mW average power at the back focal plane of the objective) controlled with an electro-optical modulator (*Conoptics*, Model 302 RM). GCaMP fluorescence was excited through a 40x water immersion objective (Nikon NIR Apo, 0.8 NA, 3.5 mm WD) and fluorescence signals detected with photomultiplier tubes (*Hamamatsu 7422P-40*), acquired with PrairieView software (*Prairie*) at 30fps frame rate (512×512 pixels, 1.3 μm/pixel). A custom dual stage preamp (1.4×10^5^ dB, Bruker) was used to amplify signals prior to digitization. Two goniometers (Edmund Optics) were used to adjust the angle of each mouse’s head in order to achieve the same imaging plane over multiple sessions.

#### Data processing for Ca^2+^ imaging

Movies were motion corrected using NoRMCorre algorithm using a non-rigid registration method that splits the field of view (FOV) into overlapping patched that are registered separately then merged by smooth interpolation^24^. Videos were then spatially and temporally down-sampled by 2 to reduce noise and the computational power required for cell segmentation. Spatial and temporal components for individual cells were extracted using large-scale sparse non-negative matrix factorization (CNMF/CaImAn)^25^ or using the singular value decomposition method by Suite2p algorithm (https://github.com/cortex-lab/Suite2P), both of which resulted in similar regions of interest (ROIs). We used Suite2p graphical user interface to manually select small, densely packed DG granule cells and discard large isolated cell bodies corresponding to mossy cells or other hilar interneurons. To obtain total number of DG granule cells within the imaging fields of view in a subset of total sessions, time averaged images were segmented using the Cellpose algorithm (https://github.com/MouseLand/cellpose) followed by manual inspection. Ca^2+^ transient events were defined by a custom detection algorithm which identifies fluorescence peaks with a rise slope greater than 4 standard deviations above an iteratively refined baseline.

#### Behavioral and Calcium Data Alignment

Behavioral data was aligned to Ca^2+^ data using the record of a synchronization signal between the two computers used for data collection. Behavioral data was down-sampled to match Ca^2+^ imaging data.

### Data Analysis

Data were analyzed using custom-written routines implemented in MATLAB. Plots were generated in MATLAB and Prism.

#### Identification of spatially-tuned neurons

We restricted our analysis to continuous running at least 2 sec in duration and with a minimum peak speed of 5 cm/sec. For each lap crossing, position data and Ca^2+^ transient events for each cell were binned into 2 cm-wide windows (100 bins), generating raw vectors for occupancy-by-position and calcium transient numbers-by-position which were then circularly smoothed with a Gaussian kernel (*SD* = 5 cm). A firing rate-by-position vector was computed by dividing the smoothed transient number vector by the smoothed occupancy vector. Within each lap, we circularly shuffled the positions 1000 times and recomputed firing rate-by-position vectors to generate a null distribution for each spatial bin. A spatially selective cell was defined that met the following criteria: (a) the cell should fire above its mean firing rate within its spatial field in at least 20% of laps or for a minimum of 3 laps; and (b) observed firing should be above 99% of the shuffled distribution for at least 5 consecutive spatial bins (10 cm) wrapping around the two edges of the belt. We have identified spatially tuned neurons by excluding bins in which sensory cues were omitted or shifted and calculated firing rate vectors in these laps separately. Among all of the spatially tuned neurons, “middle cue cells” were defined as those with averaged spatial fields that overlapped with at least 50% of the 45^th^ and 55^th^ bins and had peak amplitude at least two times larger than those in cue-omitted laps. “Lap-cue cells” were defined as those with averaged spatial fields overlapping at least 50% of the region wrapping around the 90^th^ and 10^th^ bins in the normal laps and have peak amplitude in cue-omitted laps and cue shifted laps not exceeding than at least two times of that in normal laps. The remaining cells constituted the “place cells”.

#### Spatial information, stability, consistency, and emergence of spatial fields

To calculate a measure for spatial information content for granule cells in Fig. 1g, we adapted a traditional method of spatial information assessment^23,38,57^ to Ca^2+^ imaging data. For each cell, we used the firing rate-by-position vector and shuffled null distribution computed above and calculated the spatial information content for each as described previously^38,57^. To account for the fact that low firing rates artificially produce high spatial information scores, we subtracted the mean of the shuffled information per spike from observed information per spike, divided by the standard deviation of the shuffled values to determine the spatial variance for each cell. Therefore, the amount of spatial information is inferred from differences in normalized Ca^2+^ activity in each neuron and reported as bits per seconds. The consistency of place field firing in Fig. 1h was determined as the cross-correlation between the averaged firing rate-by-position vector of the first and the second halves of the total number of cue normal laps within a session. We determined place field onset lap in cue normal laps (Fig. 1i) as described previously^39^. Briefly, starting on lap 1 we searched for a significant Ca^2+^ transient event present within the boundaries of the previously determined mean spatial field calculated from all the laps in the session. If one were found we would then search for Ca^2+^ transient event on each of the next 4 laps. If 3 of the 5 laps had Ca^2+^ transients within the mean place field boundaries, lap 1 would be considered the place field onset lap. If either lap 1 had no Ca^2+^ transient or less than 3 of the 5 laps had Ca^2+^ transient, we would move to lap 2 and repeat the search (Fig. 1i). The field onset laps, and the mean firing rates during first 5, 6-10 and 11-15 laps was then assessed for each group of cells.

#### Multi-Session Cell Tracking

Cells were tracked across sessions using CellReg^28^. Briefly, rigid alignment with both translations and rotations was performed on spatial footprint projections of each session and manually inspected for quality. To improve performance with our data, we modified the CellReg source code to consider complete spatial footprints instead of centroids during alignment. The centroid distance between neighbors was then calculated and used to create a probabilistic model that estimated the expected error rate at different thresholds. The optimal centroid distance threshold was chosen by the algorithm and used to match cells. A clustering algorithm then refined these decisions previously made using pairwise comparisons.

Following cell registration, tracked cells were matched with their corresponding functional cell types (i.e. mid-, lap-cue, place cells, as described above). All analyses presented in Figure 3 are carried out in pairwise, to maximize the number of cells in each comparison and to minimize the total number of comparisons. For multiday comparisons we used Day1, session 1 as the normal session, and for multisession comparisons we used Visual stimulus session as the normal session. To calculate the fraction of cells that maintain their identity, cell pairs that were counted as being the same cell type in both sessions was divided by all of that cell type in the normal session. In order to derive a null distribution for preservation of pairwise identity, we randomly permutated the cell IDs of all the tracked cells in pairwise sessions 1000 times and calculated the fraction of cells that were the same, among all of that cell type in the normal session. We calculated p-values by comparing actual data to this null distribution, 97.5^th^% of the null distribution is presented dotted lines in Figure 3.

#### Rate correlation and population vector (PV) analysis

Comparison of the activity between different sessions was calculated using Pearson’s correlation of the spatially binned, averaged firing rate-by-position vector in cue normal laps in Figures 2,3, S4. The variability in neural activity between self-driven and motorized treadmill sessions (Figure 2) and between lap types (in Figure 6) was calculated by using Pearson’s correlation on each 2 cm bins of the firing rate-by-position vector along the treadmill during odor cue trials (mean for all spatially tuned cells).

#### Spatial modulation

To compare single cell cue responses at different locations, cue-triggered average z-scored calcium transients were measured for cue cells, defined with reference to omit lap activity as done previously, for middle cue sessions. For dual cue location experiments, cue cells were identified as units with receptive field peaks in the cue location whose cue-zone calcium event rate was greater than 95% of shuffled responses versus the cue-omitted laps. For statistics of spatial modulation over days, cue cells that significantly preferred one cue location were counted when responses at one location exceeded 95% of the shuffled cue position event rates. Spatial modulation in this paradigm is listed as a ratio of mean spatial firing rates for the non-preferred location over the preferred location rates for each cell (i.e. lower values correspond to higher spatial selectivity of cue responses).

#### Bayesian Reconstruction Analysis

To calculate the probability of the animal’s position given a short time window of neural activity, we used a previously published method based on Bayesian reconstruction algorithm^27^. Briefly, Ca^2+^ transient events for each cell were binned into 1 second windows to construct firing rate vectors. For each of these binned firing rate vectors, Bayesian classification of virtual position (posterior probability for each bin) was performed by a previously described method^27^ utilizing a template comprising of a cell’s smoothed firing rate-by-position vectors. In order to cross-validate our decoding procedure, we divided firing rate-by-position template into lap crossings, used 1/5^th^ of laps as “testing” dataset while the remaining 4/5^th^ of laps constituted the “training” dataset. For example, lap 1 was tested based on the firing rate-by-position vectors calculated using laps 2,3,4,5, lap as template, and lap 6 was tested based on the firing rate-by-position vectors calculated from laps 7,8,9,10, and so on. The resulting posterior probability distribution for each bin is the likelihood for an animal is located in that bin, which adds up to 1, and the bin with the maximum posterior probability is the estimated position of the animal. To determine the decoding error we calculated the absolute difference between the animal’s actual position and the maximum posterior probability in that bin. Post-reconstruction, we divided the time bins (excluding those with no activity) according to the lap types.

#### Single cell cue shift analysis

Spatial firing rates for each spatially tuned cell on normal middle cue laps were cross-correlated with firing rates on shift laps in order to estimate the cross-correlation peak offset (i.e. shift magnitude) for each cell after cue manipulation. Binned histograms of numbers of cells with spatial receptive fields at particular locations along the track were plotted with respect to their shift magnitudes. Numbers of cells were averaged over populations showing no shift (−5 to 5 cm shift) or that shifted their firing along with the cue (50cm±5cm).

#### Inhibition analysis

Out-of-field firing was calculated for cells found to be significantly spatially tuned on normal middle cue laps by extracting calcium event rates in the ~200cm track length excluding the peak place field (+/− 10cm). Average out-of-field firing rates were then calculated across all cells for cue laps and intermittent laps where the cue was omitted. For comparison of cue-associated excitation and inhibition levels, average firing rates were computed by session for the 20cm region surrounding the middle cue, with respect to the normal pre-cue baseline firing rate. Cue-related inhibition of place cell firing was calculated by selecting spatially tuned cells whose firing field on normal middle cue laps fell within the region of the cue on shift laps (50-80cm). Firing rates for these cells were then averaged for normal laps where the cue was not presented in this region and compared with laps where the cue was shifted to this region (50cm). Mutual inhibition between cues was calculated by first selecting cells responsive to each of 2 cues of different modalities (olfactory or visual) presented at 40cm and 120cm. Responses were then averaged for each cue cell for laps in which the cues were presented alone at these locations versus intermittent laps where the cues were presented together at one of the two former locations.

### Data and Software Availability

Data and custom programs are available upon reasonable request.

## Supplementary Figures

**Supplementary Figure 1.**
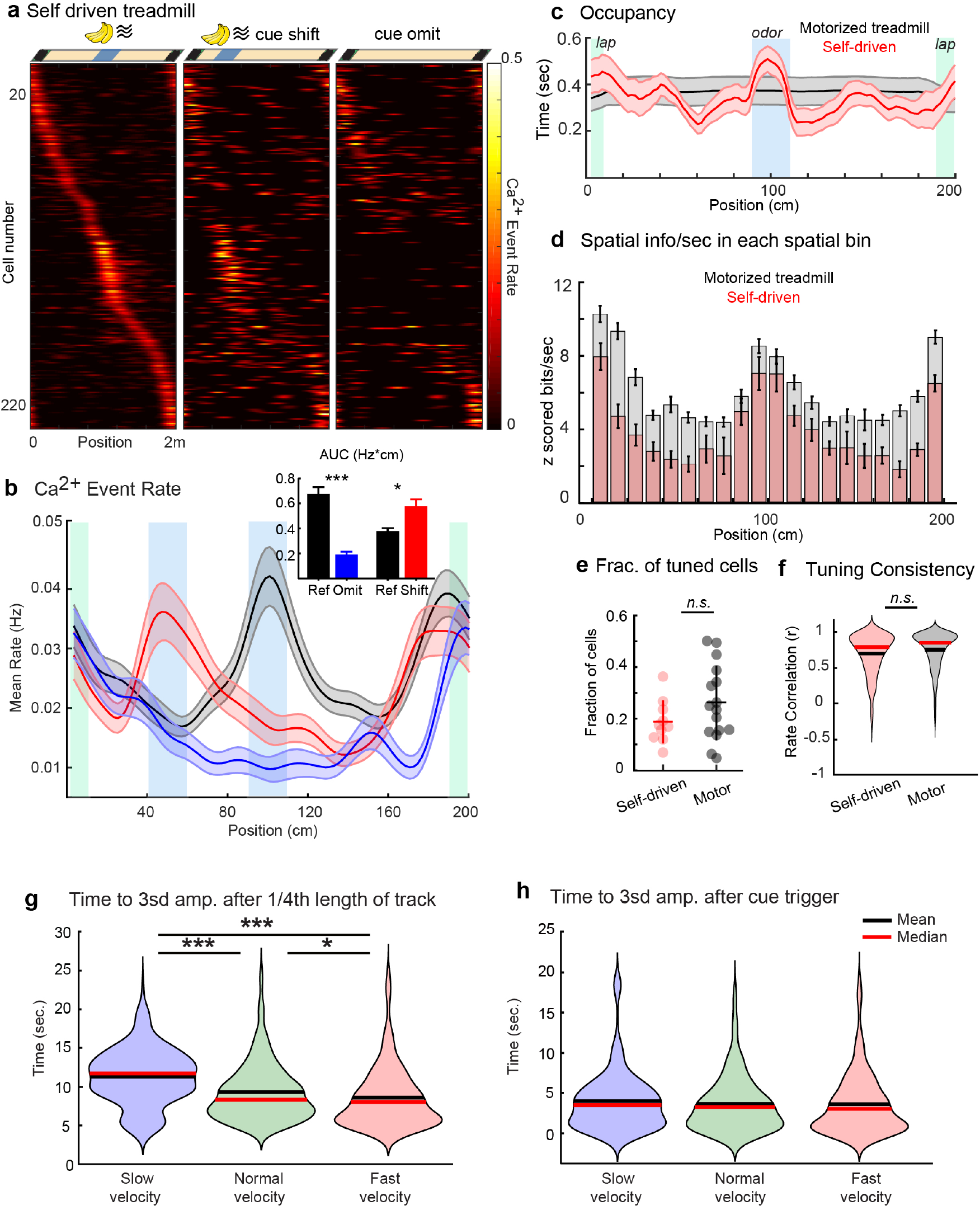
Cue associated activity and spatial properties of DG neurons are similar in self-driven and motorized treadmills, and at different treadmill motor velocity. **a)** Spatial firing patterns of DG neurons during self-driven locomotion in the treadmill spatial cue task. (Left) normal middle cue lap responses, (middle) cue shift laps, (right) laps in which odor cue is omitted (n=234 cells from 10 sessions in 5 mice). Cells sorted by activity on normal middle cue laps. **b)** Average spatial firing rates from all cells in “a.” during normal middle cue (black), cue shift (red), and cue omitted laps (blue). (Inset) averaged area under the firing rate curves (Hz*cm) within the middle and shift location reference regions during normal (black bars), cue omitted (blue bar), and cue shifted laps (red bar). *P_Normal-Omit_* <0.001, *P_Normal-Shift_* =0.03, Wilcoxon signed rank test, error bars are mean ± SEM. Each row across all graphs represents a single cell from the same session. **c)** Spatial occupancy of each mouse is calculated as the time spent in each 2cm bin along the 2 meter long treadmill during self-driven (red) or motorized (black) locomotion. Occupancy is shown for normal middle cue laps. During self-driven sessions mice tend to slow down upon odor cue delivery, which we aimed to normalize by using motorized treadmills. **d)** Mean spatial information by position showing similar enrichment of neurons with high spatial information around the sensory cues in both self-driven (pink) and motorized (gray) treadmill running. Average Z scored spatial information for cells binned by tuning position during the spatial cue task performed on a motorized vs. self-driven treadmill. Error bars represent ± SEM. **e)** Fraction of active cells that are spatially tuned (with at least 0.001 transients per s) in mice advancing the treadmill belt through self-driven locomotion (5 mice, 2 session/mouse) and mice running on the motorized treadmill (8 mice, 2 session/mouse, p=0.4508 Wilcoxon Rank Sum test) **f)** Tuning consistency of cue and place cells during self-driven vs. motorized treadmill running. Firing rate correlation between first and last halves of the session, p=0.3197 Wilcoxon Rank Sum test. **g)** Variable speed motorized treadmill: Averaged latencies of *reference position-triggered* Ca^2+^ transients of odor cue cells during running on treadmill motorized at fast, normal and slow velocities, χ^2^=85.19, p<0.001, *P_Fast-Normal_*<0.0001, *P_Fast-Slow_*<0.0001, *P_Normal-Slow_*=0.0295, Friedman test and Dunn’s multiple comparisons tests. **h)** Averaged latencies of *cue triggered* Ca^2+^ transients of same odor cue cells in “g.” during running on treadmill motorized at fast, normal and slow velocities, χ^2^=4.99, p=0.0823, Friedman test. In **(g)** and **(h)**, n=25 odor cue cells, 3 mice, red lines in violin plots show median, black lines show mean.

**Supplementary Figure 2.**
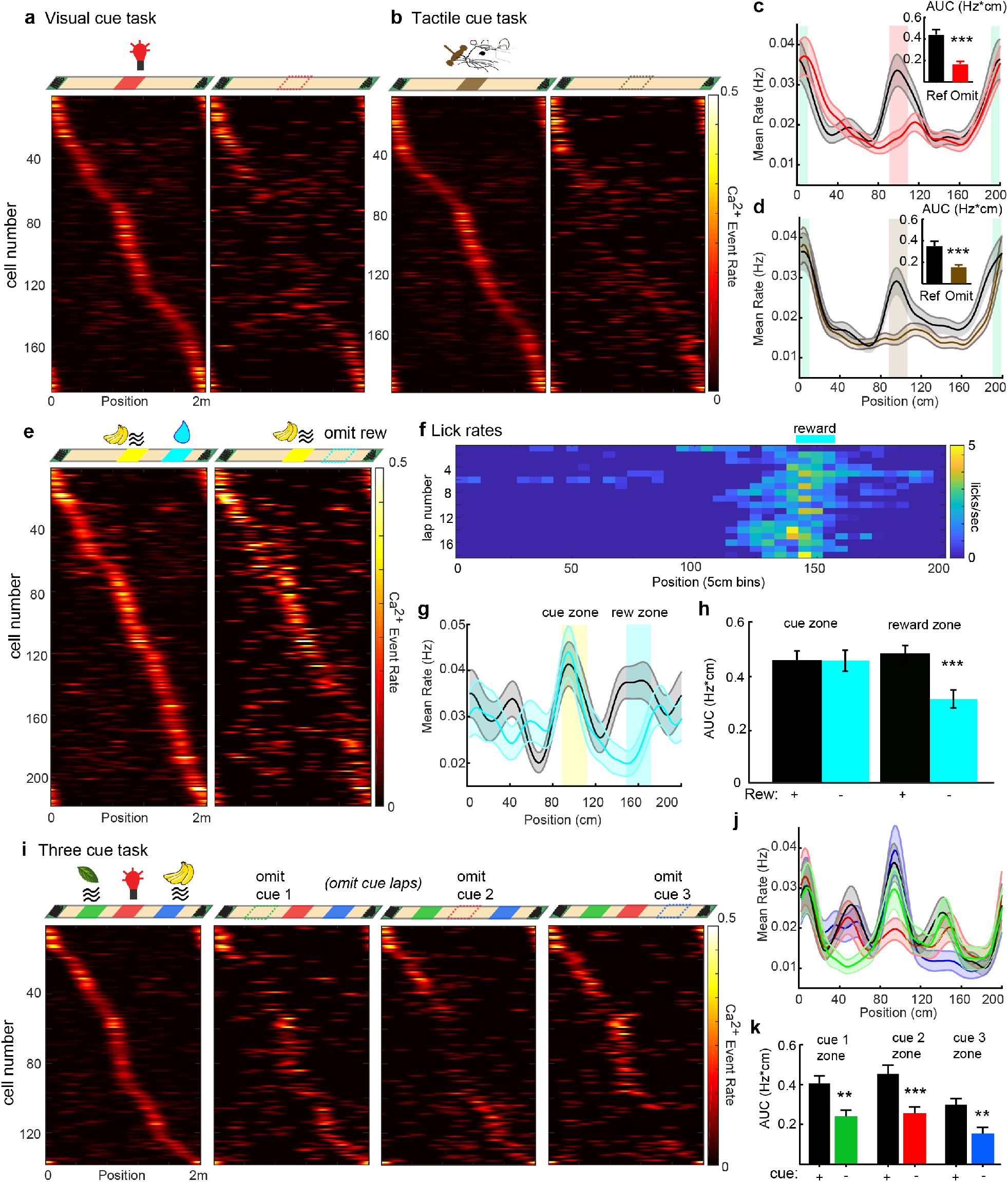
Visual and tactile cues, reward and sequences of cues evoke robust responses in DG. **a)** Different modality cues: Firing of DG neurons in response to an LED visual cue. Top: Location of lap cue (green boxes) and an LED cue (red box), during normal and cue-omitted laps. Bottom: spatial firing rates of 188 spatially tuned neurons (n=6 mice) during the first session of exposure to the middle visual cue on normal (left), and cue-omitted (right) laps. **b)** Firing of DG neurons in response to a whisker tactile cue. Top: Location of lap cue (green boxes) and a whisker tactile cue (a piece of rough sandpaper, brown box), during normal and cue omit laps. Bottom: spatial event rates of 199 spatially tuned neurons (n=8 mice) during the first session of exposure to the tactile cue on normal (left), and cue-omitted (right) laps. Each row represents an individual cell, and cells are sorted by their activity in middle cue laps. **c)** Average spatial event rate for neurons shown in “a.” (LED cue) on normal (black, mean ± SEM), and cue-omitted (blue) laps. Inset shows the averaged area under the firing rate curves (AUC, Hz*cm) within the visual cue region during normal laps (black bar) compared to the same region during cue omitted laps (blue, p<0.0001, Wilcoxon Signed Rank Sum test). **d)** Average event rates of neurons shown in “b.” (tactile cue) on normal (black), and cue-omitted (blue) laps. Inset shows the averaged area under the firing rate curves (AUC, Hz*cm) within the tactile cue region during normal laps (black) compared to the same region during cue omitted laps (brown, p<0.0001, Wilcoxon Signed Rank Sum test). **e)** DG reward responses: Top: Behavioral task with a static operant reward at 75% (150cm) track length, preceded by an olfactory cue in middle of track. Bottom: Spatial event rates for all tuned cells in the static reward task. Laps with olfactory cue in middle of track and reward at 150cm (static rew., left), or laps with reward omitted (omit rew., right). (n=256 cells, 4 mice, 7 sessions). **f)** Averaged spatial lick rates over laps, for behavioral sessions in “e.”. **g)** Average spatial event rates over all cells in “e.” during static reward laps (black), and omit reward laps (blue). Olfactory cue zone shaded in yellow, reward zone shaded in blue. **h)** Averaged area under the firing rate curves (AUC, Hz*cm) for cue zone and reward zone on static reward (black) and omit reward laps (blue). p = 3.8e^−9^, reward zone rate during rew+ vs. rew-(omit rew.) laps, Wilcoxon signed rank test. **i)** Cue sequence task: (Top, left) Experimental setup with 3 cues, cue 1 = mint odor (olfactory), cue 2 = LED (visual), cue 3 = isoamyl acetate odor (olfactory). Individual cues are omitted on intermittent laps (right). (Bottom) Spatial event rates for the same tuned cells in normal 3-cue laps (left), and intermittent laps where individual cues are omitted (right 3 columns, as labeled, n=138 cells from 5 mice, 1 session/mouse, sorted by normal 3-cue lap activity). **j)** Mean spatial event rates over all cells in “e.” on normal laps (black), and laps where cue 1 is omitted (green), cue 2 omitted (red), or cue 3 omitted (blue). **k)** Averaged area under the event rate curves (AUC, Hz*cm) within zones corresponding to cues 1-3 on normal laps (black), laps where cue 1 is omitted (green, p= 0.0031), cue 2 omitted (red, p<0.001), or cue 3 omitted (blue, p=0.0047). Wilcoxon signed rank test, error bars are mean ± SEM.

**Supplementary Figure 3.**
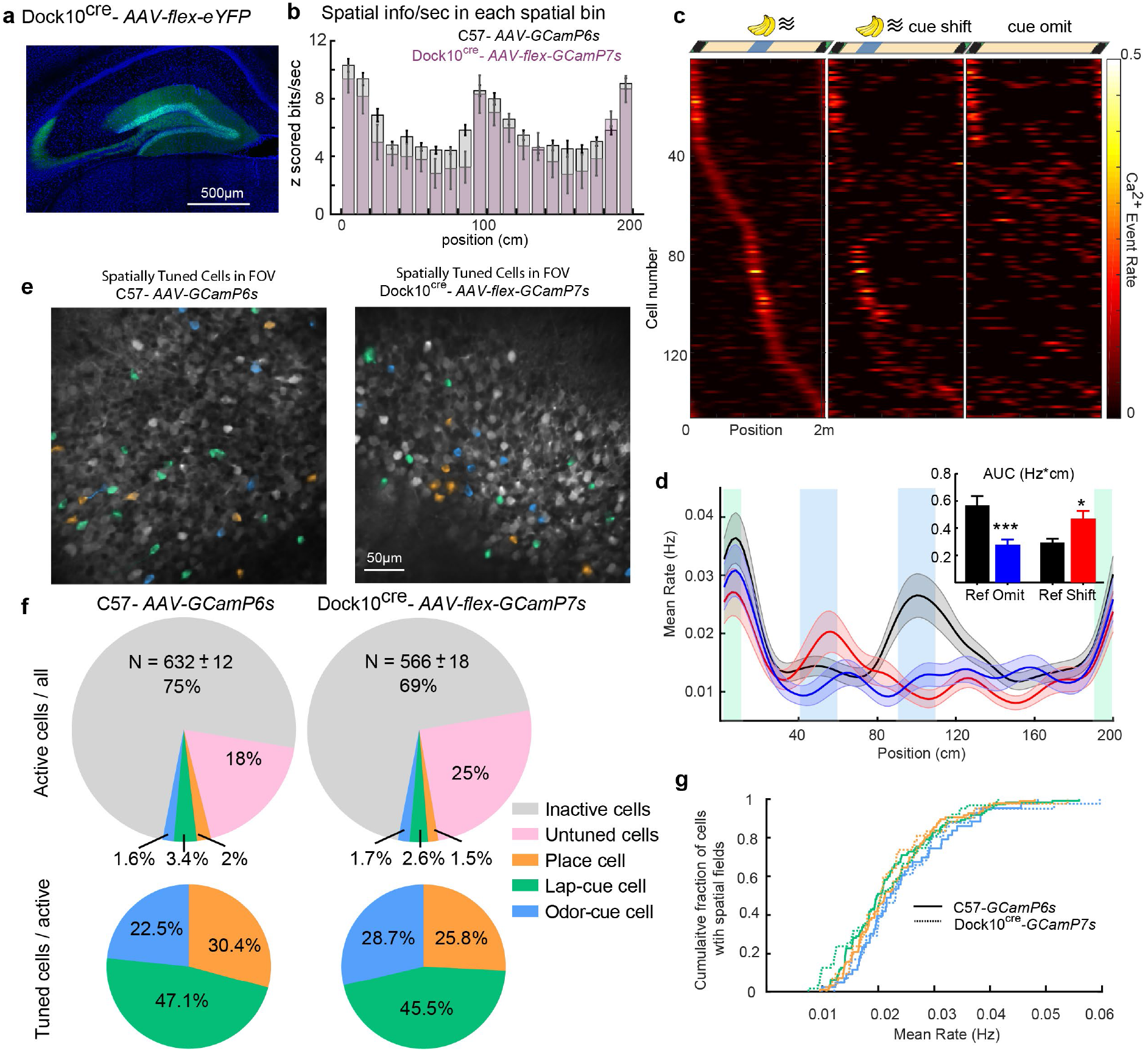
Representations of sensory cues and place in a granule cell specific transgenic mouse, Dock10-cre. **a)** Confocal image projection of brain slice from Dock10-cre mouse injected with FLEX-eYFP AAV in the dorsal dentate gyrus, showing selective labeling of granule cells (blue=DAPI, green=eYFP). **b)** Mean spatial information by position in Dock10-cre mice. Average Z scored spatial information for cells binned by tuning position during spatial cue task. WT dorsal dentate gyrus neurons express GCamP6s (gray bars) and Dock10-cre labelled granule cells express FLEX-GcamP7s (purple bars). Error bars represent SEM, *P_spkZ-DG–spkZ-GC_* = 0.15, Wilcoxon rank-sum test calculated over cumulative mean spkZ. **c)** Spatial firing rates of Dock10/GCaMP7s-expressing granule cells in the spatial cue task with an olfactory cue (IAA/banana). (Left) normal middle cue responses, (middle) cue shift laps, (right) laps in which odor cue is omitted (n=146 cells from 4 sessions in 2 mice). Cells sorted by activity on normal middle cue laps **d)** Average spatial firing rates from all cells in “c.” during normal middle cue (black), cue shift (red), and cue omitted laps (blue). (Inset) Averaged area under the event rate curves (AUC, Hz*cm) at the middle cue location during normal (black bar) and cue omitted laps (blue bar), and at the shifted cue location during normal (black bar) and cue shifted laps (red bar). *P_Normal-Omit_* <0.001, *P_Normal-Shift_* =0.03, Wilcoxon matched-pairs signed rank test, error bars are mean ± SEM. **e)** A representative *in vivo* two-photon imaging field of view within the dentate gyrus of C57 mice expressing GCaMP6s (left) and Dock10-cre mouse expressing FLEX-GCaMP7s in granule cells (right), including spatially scattered odor (blue) and lap (green) cue cells and place cells (orange). **f)** Fraction of inactive cells (gray), active cells (with at least 0.001 transients per s, pink) and spatially tuned cells (orange = place cells, green = lap cue cells, blue = odor cue cells). Mean_C57-GcamP6s_ = 631.68±11.9 cells, from 8 mice 2 sessions each, Mean_Dock10Cre-GcamP7s_ = 566.11±18.4 cells, from 2 mice 2 sessions each. Small, round somata corresponding to granule cells within the granule cell layer were counted using the *Cellpose* algorithm followed by manual inspection in a subset of total sessions imaged from one mouse. **g)** Cumulative distribution curve of the firing rate of odor cue, lap cue and place cells in Dock10-cre mice (thin lines) vs. C57 mice (thick lines); Odor Cue *P_C57-GcamP6s-Dock10Cre-GcamP7s_*=0.11 (blue), Lap Cue *P_C57-GcamP6s-Dock10Cre-GcamP7s_* = 0.05 (green), Place cell *P_C57-GcamP6s-Dock10Cre-GcamP7s_* = 0.05 (orange), Comparisons are Wilcoxon rank-sum test.

**Supplementary Figure 4.**
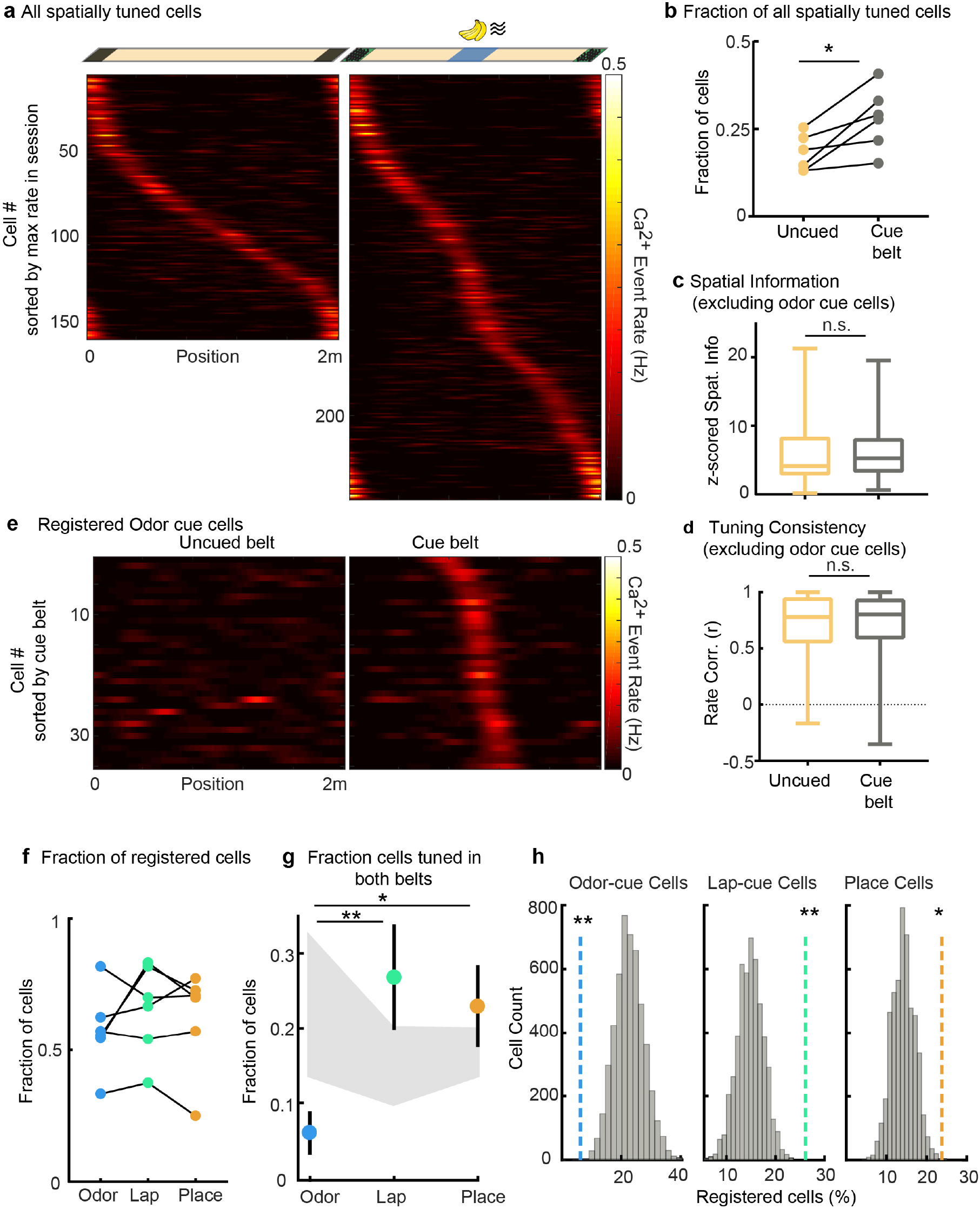
Odor cue cells do not emerge from place cells. **a-c)** Comparison of cell tuning properties between uncued vs. cued treadmill belts. **a)** Spatial firing rates of all tuned DG neurons within the same fields of view (FOV) recorded during paired sessions on an uncued belt (left) and subsequently an odor cued belt (right). Neurons are ordered according to the position of their spatial receptive field in each belt (uncued: 155, cue: 249 spatially tuned neurons in 6 mice, 1 session/mouse), but are not matched between uncued and cue plots. **b)** Fraction of active cells that are spatially tuned (with at least 0.001 transients per s) in matched FOVs recorded on uncued and cue belts (p=0.0313, Wilcoxon signed-rank test, n=6). **c)** Spatial information of all spatially tuned neurons, excluding odor-cue responsive cells, in uncued (n=155) and cued sessions (n=193, p=0.0897, Wilcoxon rank-sum test). **d)** Tuning consistency (firing rate correlation between first and last halves of the session) of all spatially tuned neurons in uncued and cued sessions, excluding odor-cue responsive neurons (p=0.5237, Wilcoxon rank-sum test). Boxes, 25th to 75th percentiles; bars, median; whiskers, 99% range. **e-h)** Comparison of individual cue cell and place cell firing between uncued and cue belts. **e)** Activity of cross-registered odor cue cells in uncued and cue belts. Cells were classified as odor-cue cells with respect to the cued belt and then their responses were determined for the uncued belt, if active during this session. Note that these “future” cue cells were not spatially tuned in uncued sessions (i.e. not place cells). **f)** Fraction of odor-cue (blue), lap-cue (green) and place cells (orange) identified on the cued belt that are cross-registered in uncued sessions (p=0.838, n=6 matched sessions, Friedman’s test). **g)** Fraction of registered cells that are spatially tuned in uncued belt sessions and encoded the odor cue (blue), lap cue (green), or place (orange) on the cued belt (χ^2^=10.88, p=0.0043, *P_OdorCue-LapCue_*=0.0066, *P_OdorCue-Place_*=0.0286, *P_LapCue-Place_*=0.9076). Gray area represent 2.5^th^ and 97.5^th^% of null distributions for each cell type. Error bars, mean ± SEM. **h)** The fraction of cross-registered odor cue cells that were spatially tuned (i.e. place cells or lap cue cells) on the uncued belt is significantly below the null distribution (left) while the fraction of lap-cue (middle) and place cells that are spatially tuned remain significantly above the null distribution. The null distributions are generated for each cell type by randomly permuting cell IDs of all cross-registered neurons and determining overlap among cell types. (Level of significance for 5,000 shufflings **p < 0.01; *p < 0.05, N_Odor-cue_=35, N_Lap-cue_=54, N_Place_=58).

**Supplementary Figure 5.**
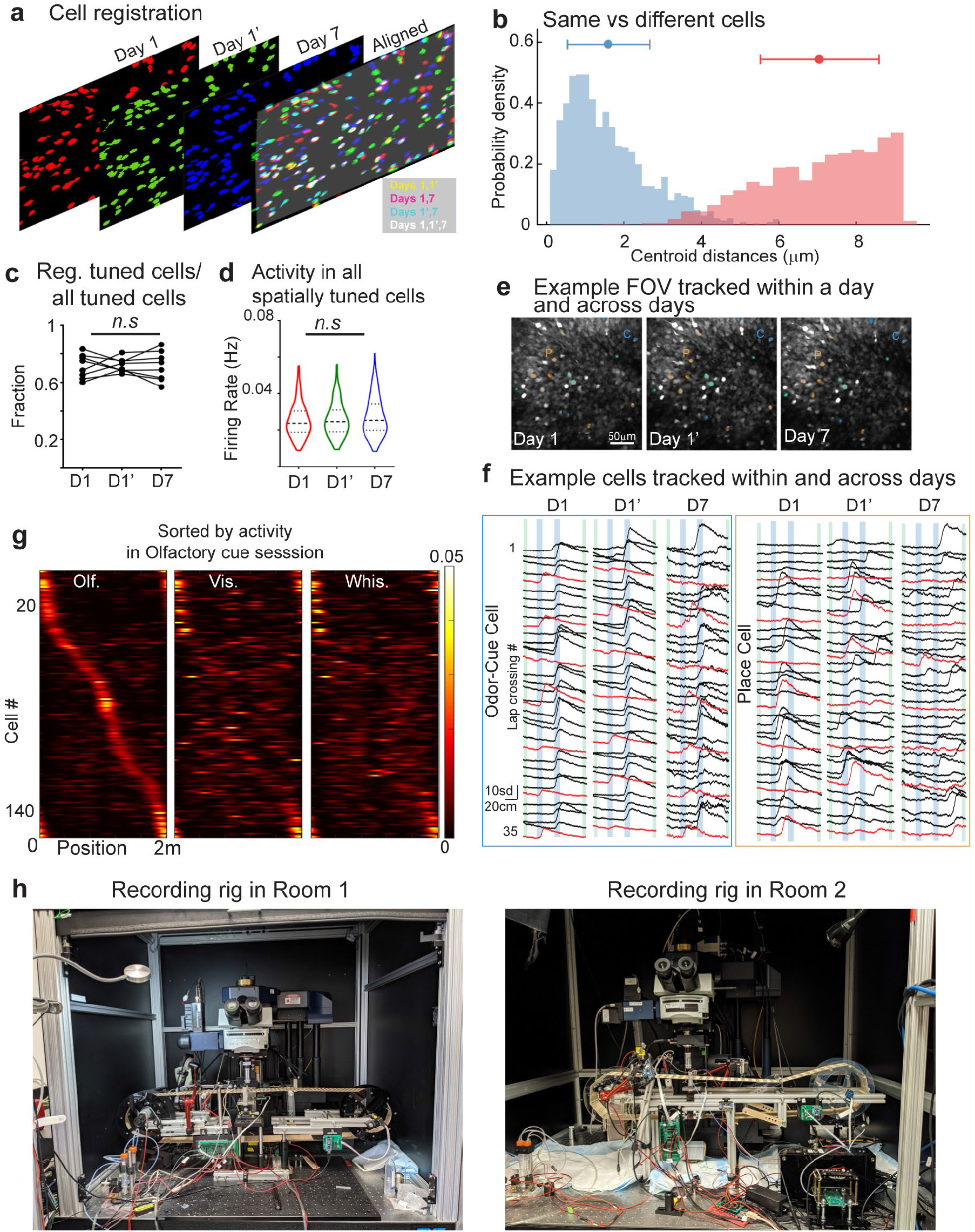
Multisession tracking of individual cue and place cells over time, with different sensory cues, and in different rooms. **a)** Representative alignment of spatial footprints for cells segmented across sessions within a day or after 1wk. in a single imaging field. **b)** Distribution of centroid distances between registered (blue, 1.45 ± 0.97, n=1604) and non-registered (pink, 7.07 ± 1.5, n=2214) neighboring cell pairs (p=6.9 10^−43^, Wilcoxon Rank Sum test). **c)** Fraction of all spatially tuned cells in day 1 session1 (D1), day 1 session 2 (D1’) and day 7 (D7) that are registered to at least one other session, χ^2^=0.25, p=0.9674, *P_D1-D1’_*>0.9999, *P_D1-D7_*>0.9999, *P_D1’-D7_*>0.9999, n=8 matched sessions, Friedman and Dunn’s multiple comparisons tests. **d)** Comparison of average firing rates in all tuned cells across days regardless of tracking χ^2^=4.993, p=0.0824, *P_D1-D1’_* >0.9999, *P_D1-D7_* =0.0781, *P_D1’-D7_* =0.4789, *N_D1_* =417, *N_D1’_* =365, *N_D7_*=338, from 8 matched sessions, Kruskal Wallis and Dunn’s multiple comparisons tests. **e)** Representative fields of view with odor cue cells (blue), lap cue cells (green), and place cells (orange) tracked within a day and across days. **f)** Representative Ca^2+^ transients for an odor cue cell that has stable cue-selective activity within one day and over 1wk (left). Transients for a place cell show relatively stable firing location within day but a reorganization of spatial selectivity across days (right). Black and red traces represent normal and cue shifted laps, respectively. **g)** Spatial firing rates of neurons tracked between olfactory, visual and whisker tactile cue sessions ordered according to the position of peak activity in olfactory cue sessions (n=157 cells, 6 mice). **h)** Photos of the two-photon behavioral setups in two different rooms, used in the room switch experiment (Fig.2f-i). Two-photon microscopes were identical and behavioral apparatuses on both rigs were similar including a similar configuration of lick port and location of odor delivery tube. However, the treadmills varied slightly in geometry and the cabinet in room 2 was larger than the setup in room1. We note that the difference in distal cues in two rig set ups were minor compared to typical arena change experiments in freely moving rodents^35,39,46^.

**Supplementary Figure 6.**
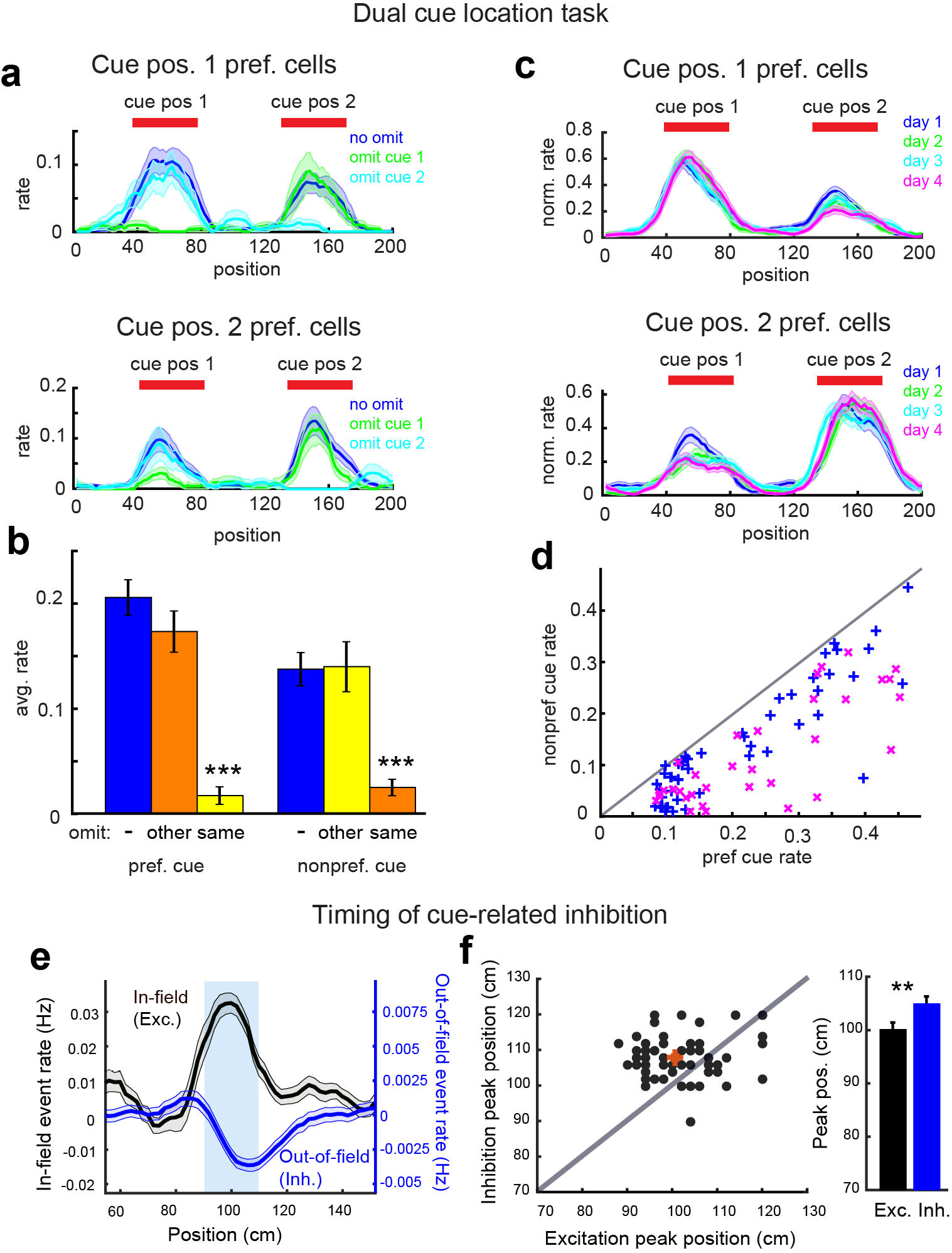
Spatial modulation of sensory cue responses and timing of cue-related inhibition in the DG. **a)** Dual cue location task: Average spatial firing rates for cue cells preferring LED visual cue position 1 (top) and cue position 2 (bottom) on normal laps (blue), laps where the cue at position 1 is omitted (green), and laps where the cue at position 2 is omitted (cyan). (n=89 pos1 cue cells, 67 pos2 cue cells). Note that cue responses for both preferred and non-preferred cue locations are largely eliminated when the respective cue is omitted. **b)** Average peak firing rates for all cue cells (both cue 1 and cue 2 preferring) for preferred cue location (left) and non-preferred location (right) on normal laps (blue), laps where the opposite cue is omitted (orange), and laps where the same cue (preferred or non-preferred) is omitted. (n=156 cells, positions1&2) (mean ± SEM: 0.205 ± 0.017, 0.173 ± 0.020, 0.017 ± 0.008, 0.137 ± 0.016, 0.140 ± 0.024, 0.025 ± 0.008, P_Preferred Cue_ = 7.16×10^−9^, P_Non-preferred Cue_ =3.48×10^−9^, Wilcoxon sign rank test). **c)** Average spatial firing rates for cue cells preferring cue position 1 (top) and cue position 2 (bottom) on days 1 (blue, n=50), 2 (green, n=40), 3 (cyan, n=50), and 4 (magenta, n=33) of dual cue location task. Rates for each cell are normalized to their receptive field center rate. **d)** Cue location firing rates for preferred and non-preferred cue locations for all cue cells on days 1 (blue, n=50) and 4 (magenta, n=33) of the dual cue location task, showing increased cue spatial preference over the recording period. **e)** Average in-field spatial firing rates (i.e. cue-related excitation) in the region around the middle cue location for all tuned neurons, compared with out-of-field, spontaneous firing rates for the same cells (i.e. cue-related inhibition), adjusted for pre-cue firing rate. Note that the position of the excitation peak precedes the trough of inhibition by 5-10cm, corresponding to 0.5-1s at typical treadmill speeds. Blue shaded area shows the cue delivery position. **f)** (Left) Session averaged excitatory (within-field) peak position vs. inhibitory (out-of-field) peak position (66 sessions). Red = avg., diagonal (gray line). (Right) Quantification of the position of excitation peak (in cm) compared to the position of inhibition peak, p<0.01, Wilcoxon Signed Rank test.

